# Decoding the constraints acting on a coastal fish using landscape transcriptomics

**DOI:** 10.1101/2025.08.05.668635

**Authors:** Emma Gairin, Saori Miura, Zoé Chamot, Camille A. Sautereau, Jann Zwahlen, Hiroki Takamiyagi, Yann Gibert, Marcela Herrera, Vincent Laudet

## Abstract

Understanding how wild organisms respond to complex mixtures of environmental stressors remains a major challenge in ecology^1,2^. To move beyond laboratory-based studies of the effect of individual stressors on model species, landscape transcriptomics – the study of how genome-wide expression patterns reveal environmental conditions – has been suggested^3^. Here, we apply landscape transcriptomics to the damselfish *Chrysiptera cyanea*, a sentinel species from Okinawa’s coastal ecosystems, by sampling juveniles and adults across 18 sites from natural to urbanized habitats. We uncover a striking molecular signature of urbanization: fish from human-impacted sites exhibit elevated expression of genes involved in inflammation and immune responses. Remarkably, their transcriptomes resemble those of overfed individuals from laboratory feeding experiments – suggesting that urban reefs act as “fast food” environments, offering energy-rich diets while imposing chronic stress. This pattern is illustrated by the altered expression of key genes in metabolic and stress pathways (*e.g.*, downregulation of *ppargc1a* and *insrb*, indicative of dietary intake surplus and insulin resistance; upregulation of interleukins, chemokines, toll-like receptors and caspase linked to the immune response and inflammation). Our results demonstrate that wild transcriptomes capture the invisible biological costs of environmental degradation, positioning fish as living biosensors for coastal ecosystem health.

## Main

Human activities alter natural habitats through local and global stressors^1^, yet current methods fall short of detecting their compounded consequences in organisms. Researchers commonly rely on parallel approaches: monitoring changes in environmental parameters, such as water quality or biological communities^4–6^ and exposing model organisms to a single or few stressors in controlled laboratory conditions to understand their impacts^7,8^. Despite their usefulness, these approaches often fail to capture the full effect of the cocktail of stressors in natural environments. With the rise of high-throughput sequencing, landscape transcriptomics – the study of how genome-wide expression patterns reflect dynamic, landscape-scale environmental drivers – has been proposed as a comprehensive method to detect multiple stressors^3,9^. Unlike targeted approaches, transcriptomics provides unbiased and hypothesis-free gene expression data^3,10–12^. Importantly, variations in gene expression can precede physiological changes, serving as an early warning signal for later-stage ecological damage such as disease outbreaks and population declines^12^. Moreover, transcriptomics can provide sensitive mechanistic insights about the consequences of environmental change^13^. Lastly, by focusing on conserved genes and pathways, transcriptomic responses can ultimately be compared across diverse taxa, enabling the construction of a common conceptual framework^14^. Landscape transcriptomics therefore shows promise as a universal tool to monitor environmental change both in wildlife and human populations. However, the application of transcriptomics at ecological scales remains in its infancy and awaits broader empirical validation.

To test the value of landscape transcriptomics under field conditions, we selected the blue damselfish *Chrysiptera cyanea,* a territorial reef species which, through its presence in a variety of shallow coastal environments, provides access to a gradient of urbanisation on the subtropical island of Okinawa, Japan^15^. These coastal zones offer contrasting conditions for reef fish; after a period of larval dispersal in the open ocean, different species may prefer specific habitat features – such as higher nutrient availability or stable temperatures – when settling on a reef^16,17^. As such, natural variability influences fish growth, survival, and potentially gene expression^18^. However, many coastal ecosystems globally are degraded due to human activities^19,20^. In Okinawa, the northern part of the island remains relatively pristine, while more than a million people live within the southern half of the 1,200 km^2^ island, with housing, industries, and agricultural activities taking place along the shore^21,22^. This is an ideal setting to study the relationship between human presence, coastal environments, and the organisms inhabiting them.

In this study, we predicted that urbanisation leads to detectable shifts in gene expression in juvenile and adult *C. cyanea*. Gene expression is however known to be influenced by factors such as sampling time, season, developmental stage, or reproductive status, which have been suggested to obscure patterns of interest in field samples^3^. To address the co-occurrence of multiple environmental factors, our sampling design drew from strategies developed for experiments with multiple factors. Indeed, the “combinatorial explosion” frequently encountered by experimentalists can be countered by assessing subsets of randomised combinations of factors, which can gather more significant results than experiments constrained to a controllable number of parameters and that may lack a key explanatory factor^7,23,24^. We adopted a semi-randomised sampling approach by targeting 18 sites along a defined gradient of human presence, and collecting five juvenile fish of varying sizes and four to five adult fish of both sexes under a range of environmental conditions (different water temperature, salinity, chlorophyll-α, and sampling time). We focused on the gene expression of the whole body of juveniles to detect organism-wide changes in gene expression, and of the liver in adults to screen long-term consequences of human activities on energy metabolism and detoxifying processes. Using this extensive dataset, we detected the impact of individual and combinations of parameters on gene expression and found that human presence profoundly influenced immunity, cell cycle regulation, stress response, and energy metabolism. Experimental validation to extract the transcriptomic signature of food availability identified a ‘fast food’ effect of urbanisation, with adults showing both a lipid-rich diet and physiological stress. Using dimensional analyses, we summarised the relative expression pattern of pre-defined groups of genes to a single value for each sample, which we visualised with radar plots; this novel approach, tailored for landscape transcriptomics, facilitated data analysis and stands as a powerful and simple way to extract a diagnosis about organismal status.

### Non-random variability in gene expression captured in transcriptomes

We sampled *Chrysiptera cyanea* across 18 coastal sites in Okinawa, Japan, along a gradient of human presence (from 2 to 94% of natural zones within 1km of each site). As a first approach to detect variations across the sites (Fig. 1a), we found that, across the 28,173 genes in the genome of *C. cyanea*^25^, 88 to 6,348 genes significantly differed in expression level in pairwise site comparisons (*e.g.,* site A *vs.* B) for the juvenile whole-body samples, and 15 to 3,763 for the adult livers (Fig. 1b). Interestingly, the number of differentially expressed genes was significantly correlated in juvenile whole-body and adult liver samples for the same pairs of sites (Fig. 1b): there was a coherent effect of local conditions at both life stages. As only one organ was considered for adults, the number of genes and range of physiological responses expressed in the samples was lower than for juvenile whole-body samples. Lastly, juveniles – likely still acclimating to their new environment – showed numerous frequently differentially expressed genes (2,410 genes in >25% of the pairwise site comparisons; Fig. 1c) while adults – which have resided on their sites for months to years – showed site-specific profiles (251 genes retrieved in >25% comparisons); this is indicative of a transition from broad acclimation responses to long-term fine-tuned mechanisms.

**Fig. 1:**
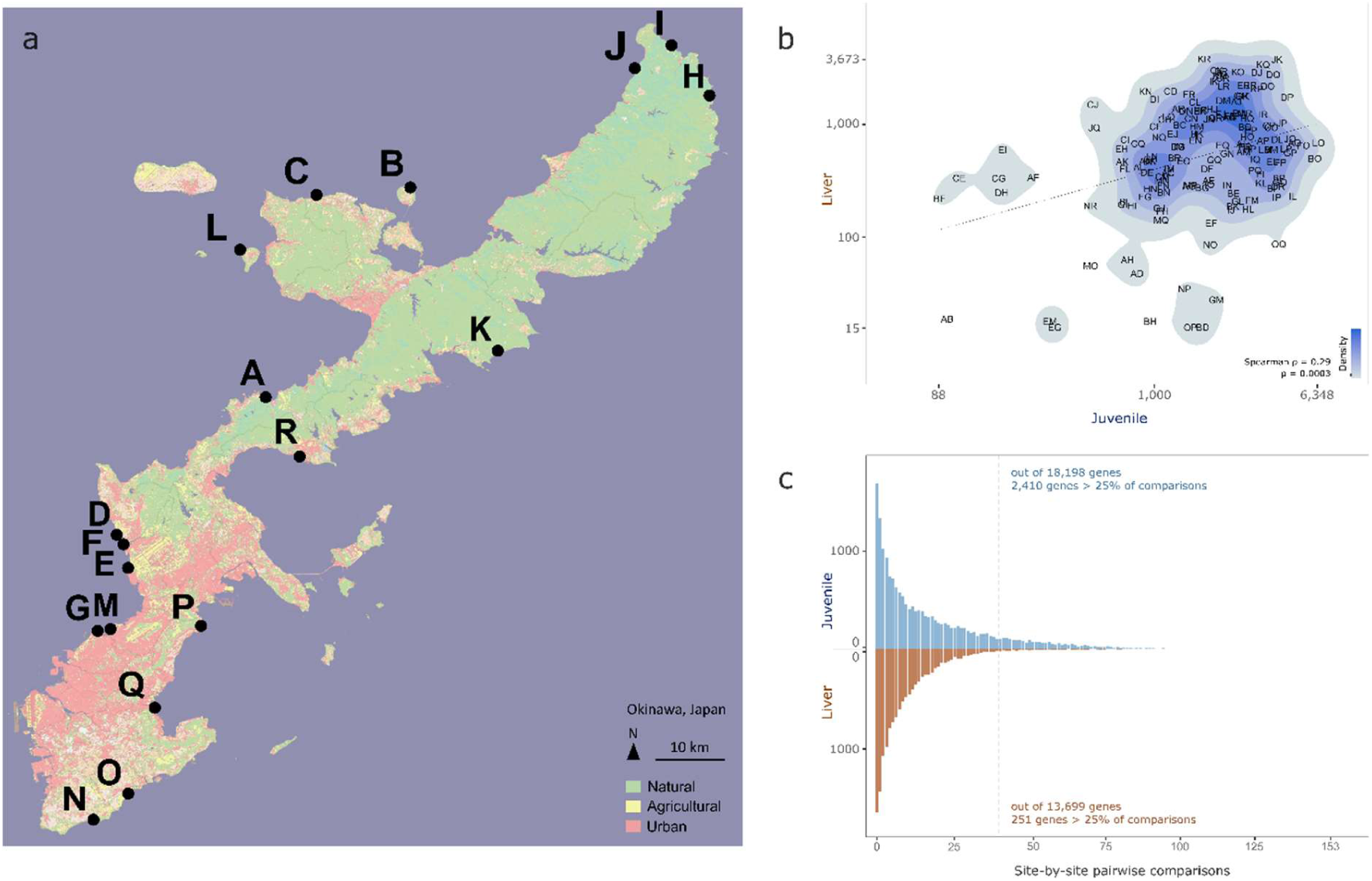
Environmental context and overview of the differential gene expression analysis for juvenile whole body and adult liver samples of *C. cyanea* across 18 coastal sites in Okinawa, Japan. **a**, Map of the eighteen field sites in Okinawa, overlaid on land-use data from JAXA (2023). **b**, Number of differentially expressed genes (logarithm-10 scale) in juvenile whole body and adult liver samples resulting from pairwise comparisons of 18 sites (absolute fold-change >1.5, p < 0.01). **c,** Number of pairwise site comparisons in which each gene was differentially expressed.

### Identifying gene expression drivers through sampling heterogeneity

Dimensional reduction using Principal Component Analyses (PCAs) showed that, although fish size and sex influenced gene expression, samples from the same sites conveyed a coherent gene expression message by clustering closely in response to local environmental drivers (Fig. 2b,c; Extended Data Fig. 1). Taking advantage of our semi-randomised sampling approach, we fitted linear and quadratic models between the expression level of each gene, the percentage of natural zones within a kilometre of the site, environmental factors (temperature, salinity, chlorophyll-α, sampling time), and fish characteristics (juveniles: standard length; adults: sex). A significant relationship with at least one of the parameters was found in >78% of the genes for juvenile whole-body and >65% for adult livers (Fig. 2d). Interestingly, 1,854 genes in juvenile whole-body and 1,312 genes in adult livers responded exclusively to temperature, which varied extensively in our sampling. When the four warmest sites (29.6-32.4°C) were excluded from the fit, the number of temperature-associated genes decreased by almost two-thirds in juvenile whole-body and by half in adult livers (Fig. 2d). Sampling time – due to circadian cycles –, fish size and sex also had a strong footprint on gene expression. A similar proportion of genes in juvenile whole-body and adult liver samples responded to the same factors, highlighting a parallel response of different tissues across developmental stages (Pearson’s Chi-squared test, χ^2^=3548, df=19, p<10^-15^).

**Fig. 2:**
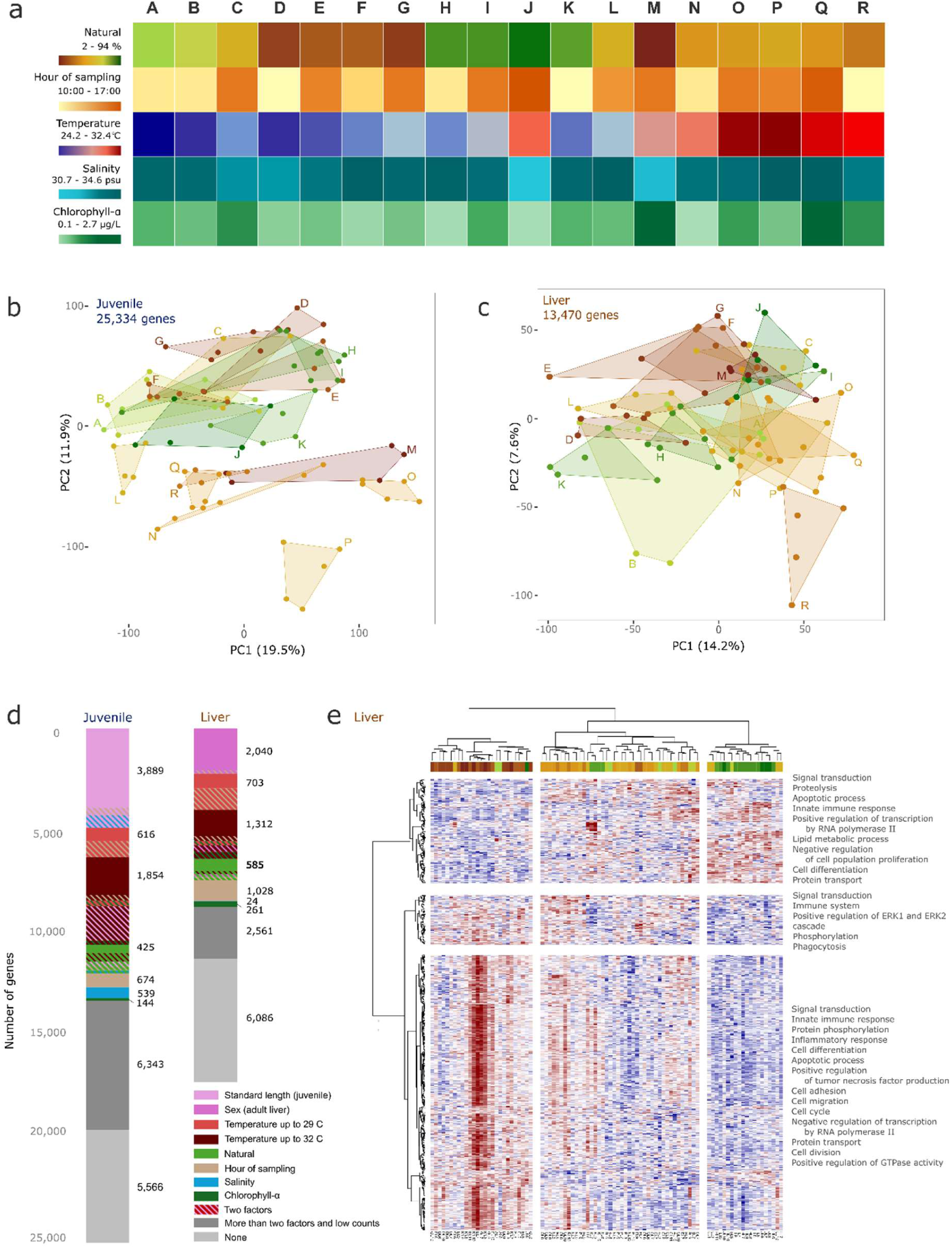
Relationship between the environment and gene expression in *Chrysiptera cyanea*. **a**, Percentage of natural zones within a kilometre of each site, sampling time, temperature, salinity, and chlorophyll-α at each site (full dataset in Supplementary Materials). **b**, Principal Component Analysis of the expression level of all genes from the transcriptomes of juvenile whole-body samples from 18 sites. Colour-coding based on the proportion of natural zones within a kilometre of the site. **c**, Principal Component Analysis of the expression level of selected genes from the transcriptomes of adult liver samples from 18 sites. 3,973 genes responding exclusively to the sex of the adult were removed from the analysis. Colour-coding based on the proportion of natural zones within a kilometre of the site. **d**, Bar plot of the number of genes with a significant linear or quadratic difference in expression level in response to fish characteristics or environmental parameters, from DRomics 2.6-2. **e,** Heatmap of the 585 genes with significant linear or quadratic differences in expression level with respect to the percentage of natural zones within one kilometre in the adult liver samples, and Gene Ontology families corresponding to the 3 main gene clusters, presented in order of prevalence within each gene cluster.

We identified 425 genes in juvenile whole-body and 585 genes in adult livers uniquely linked to human activities. These conveyed generalised stress at multiple biological levels, with modifications of the expression level of genes involved in apoptosis, immunity, and signal transduction (genes such as interleukins, chemokines, caspases; KEGG pathways including antigen processing and presentation, virus infection, T cell, toll-like, and NOD-like receptor signalling, chemokine and NF-kappa B signalling). We also found a decrease in the expression of genes related to cell differentiation, tissue homeostasis, protein transport and proteolysis, and transcription in favour of inflammation, carcinogenesis, reactive oxygen species, protein phosphorylation, and cell cycle regulation near human activities (Fig. 2e). These genes are potential markers of human-related stress for environmental monitoring in *C. cyanea* as they do not respond to other environmental or organismal factors.

### Integrating experimental and field transcriptomes to infer feeding status

Landscape transcriptomes can be assessed through the lens of local parameters (Fig. 2) but can also be compared to reference datasets from controlled conditions^26^. To test the value of combined field and laboratory approaches, we explored the impact of food availability on gene expression in juvenile whole-body and adult liver samples using two conditions: fasted or fed. Following 3-day and 14-day experiments for juveniles and adults respectively (durations tailored to their tolerance to fasting), PCAs showed a clear separation of the fasted and fed transcriptome samples along PC1 (Extended Data Fig. 2). The top 10% of genes driving PC1 in juvenile whole-body and adult liver transcriptomes were related to cell cycle and division, DNA transcription, protein modification, lipid and carbohydrate metabolism, and glucose homeostasis (Extended Data Fig. 3). We used these genes to generate secondary PCAs grouping experimental and field samples (Fig. 3a,b), from which we extracted the feeding status of each field sample using its PC1 coordinate as a proxy.

**Fig. 3:**
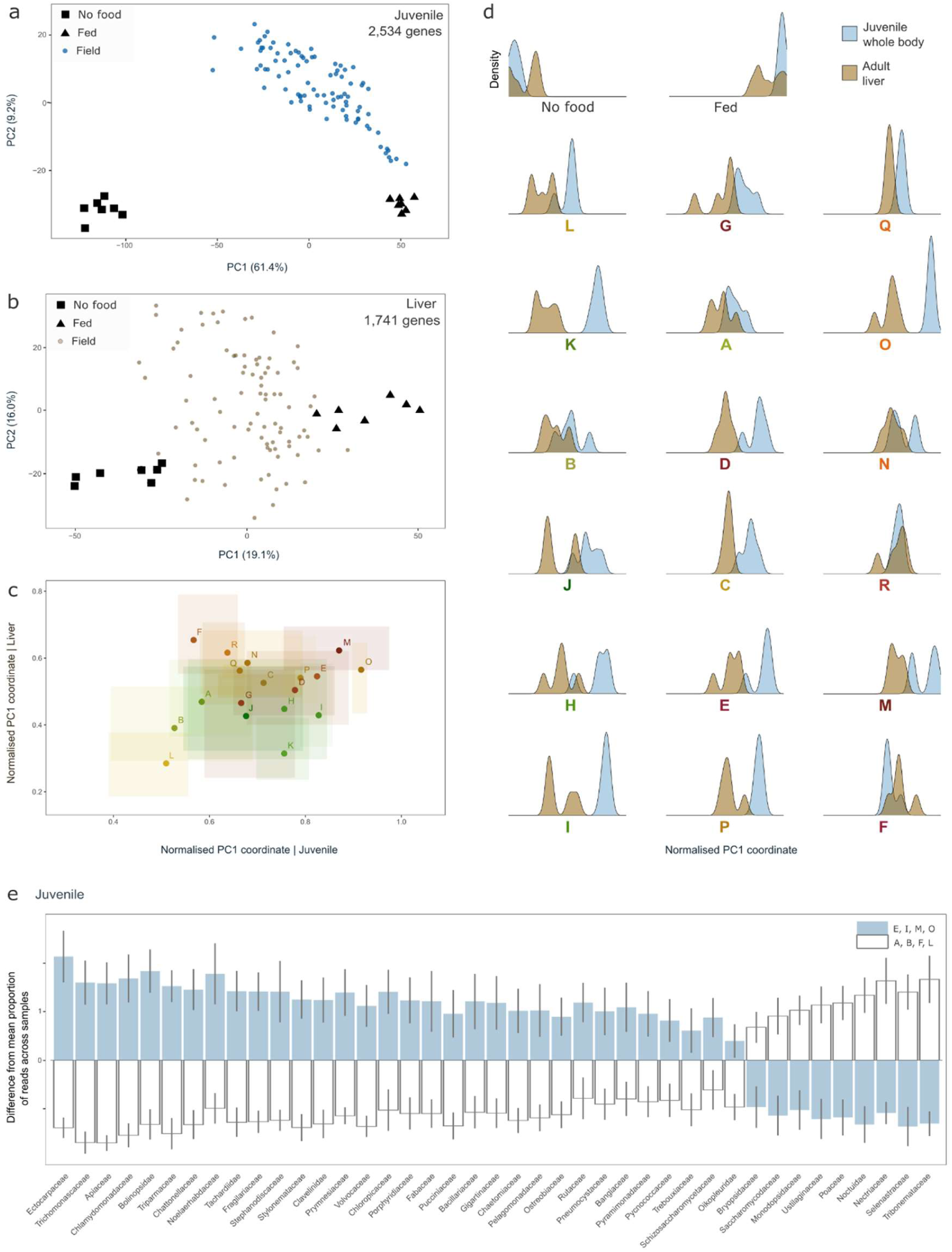
Feeding status of juvenile and adult fish from the field through a comparison with a fasted-fed laboratory experiment, and meta-transcriptomic juvenile diet analysis. **a**, PCA of the expression level of 2,534 genes in the field and laboratory experiment samples, for juvenile whole-body samples. These 2,534 genes are the top 10% contributors to the PC1 variance of a preliminary PCA using the laboratory samples. **b**, PCA of the expression level of 1,741 genes in the field and laboratory experiment samples, for adult liver samples. These 1,741 genes are the top 10% contributors to the PC1 variance of a preliminary PCA using the laboratory samples. **c**, Mean normalised PC1 coordinate of the juvenile and liver samples from each site (dot) as well as minimum and maximum coordinates (shaded rectangles). The colour coding corresponds to the percentage of natural zones within a kilometre of each field site. **d**, Density spectra of the juvenile and liver normalised PC1 coordinate on each site; sites presented in order of the mean PC1 coordinate for the adult liver samples, from low to high. **e**, Density spectra of the juvenile and liver normalised PC1 coordinate on each site; sites presented in order of the mean PC1 coordinate for the adult liver samples, from low to high. **e**, Proportion of taxonomically-assigned reads in juvenile samples deviating from the mean in the sites with highest (E, I, M, O) and lowest (A, B, F, L) feeding status.

Strikingly, adults showed higher feeding status near urban zones (linear mixed-effect model, site as a random effect, p=0.008; Fig. 3c). Among diet-related genes with a modified expression in livers from urban zones, we found Insulin Receptor b *insrb* and Peroxisome Proliferator Activated Receptor Gamma Coactivator 1 Alpha *ppargc1a*, markers of insulin resistance and energy surplus. Coupled with an upregulated inflammation and immune response (Fig. 2e), this suggests that human activities lead to “fast-food-like” habitats, where macronutrients are plentiful but fish are exposed to immune challenges. A similar trade-off was reported in transcriptomic studies of two bird species, where an upregulation of the immune response was coupled with a lipid-rich diet in urban zones^27,28^. This potential commonality in transcriptome signatures in response to human activities must be explored in more ecosystems and taxa.

In juveniles, size was positively linked to feeding status (linear mixed-effect model, site as random effect, p < 0.001). Although we found significant differences in PC1 coordinates between 39 pairs of sites (Kruskal-Wallis χ^2^=65.7, df=17, p<10^-6^ followed by Dunn test), none of the environmental parameters measured in the study was significant. We thus analysed juvenile diet using a meta-transcriptomic approach; by taxonomically classifying the sequencing reads which did not align to *C. cyanea*’s reference transcriptome, we retrieved significantly more tunicates (*Clavelinidae, Oikopleuridae*), ctenophores (*Bolinopsidae*), brown algae (*Ectocarpaceae*), and red algae (*Gigartinaceae*) reads in juveniles from sites associated with a higher feeding status (Fig. 3e).

Lastly, the mean PC1 coordinate was greater (higher feeding status) for juveniles than adults, with the exception of sites F and R (Fig. 3d); the fasted treatment could have had a stronger impact on the juveniles over three days than on the adults over two weeks, leading to a more extreme response in the fasted juvenile group. Alternatively, Okinawa’s coastal zones may provide more adequate food to juveniles than to adults.

### Radar plots link transcriptomes to local stressors

Okinawa’s *C. cyanea* transcriptomes revealed extensive differences linked to human presence, environmental parameters, and feeding (Figs. 1,2,3): they can serve as indicators of fish health and local conditions. However, the use of landscape transcriptomics by environmental managers may be impeded by the complexity of grouping and analysing numerous samples, environmental conditions, and biological processes. To tackle this multidimensionality, novel data summarisation and visualisation tools are required. Building upon manually curated lists of genes related to energy metabolism, neurotransmission, endocrine regulation, and stress response^14^, we generated PCAs for each gene group, calculated the average coordinates of each site for the top PC dimensions, and identified the axis of maximum variance; we then projected the coordinates of each sample onto this axis and set lower values to correspond to samples with overall lower gene expression (Table S7; Extended Data Fig. 5). This yielded a single value per sample and gene group representative of the position of the sample compared to all others. We then visualised these values on radar plots, enabling the comparison of samples and sites and identification of co-occurring processes and site-specific stress signatures.

These radar plot profiles were consistent in fish exposed to similar conditions, and intra-site variability could be attributed to size or sex in many cases (Fig. 4). Fish from similar sites showed closely related profiles (*e.g.,* for juveniles, A, B, and L with medium urbanisation and low food availability, N and R with similar urbanisation and water temperature; for adults, H and I, among the most natural, as well as E, G, and M, among the most urbanised). Genes related to a single and strong environmental driver directly relayed local characteristics, such as heat-shock markers in fish exposed to >31.9°C (O and P) or the bimodal phototransduction signature in juveniles depending on morning or afternoon sample collection.

**Fig. 4:**
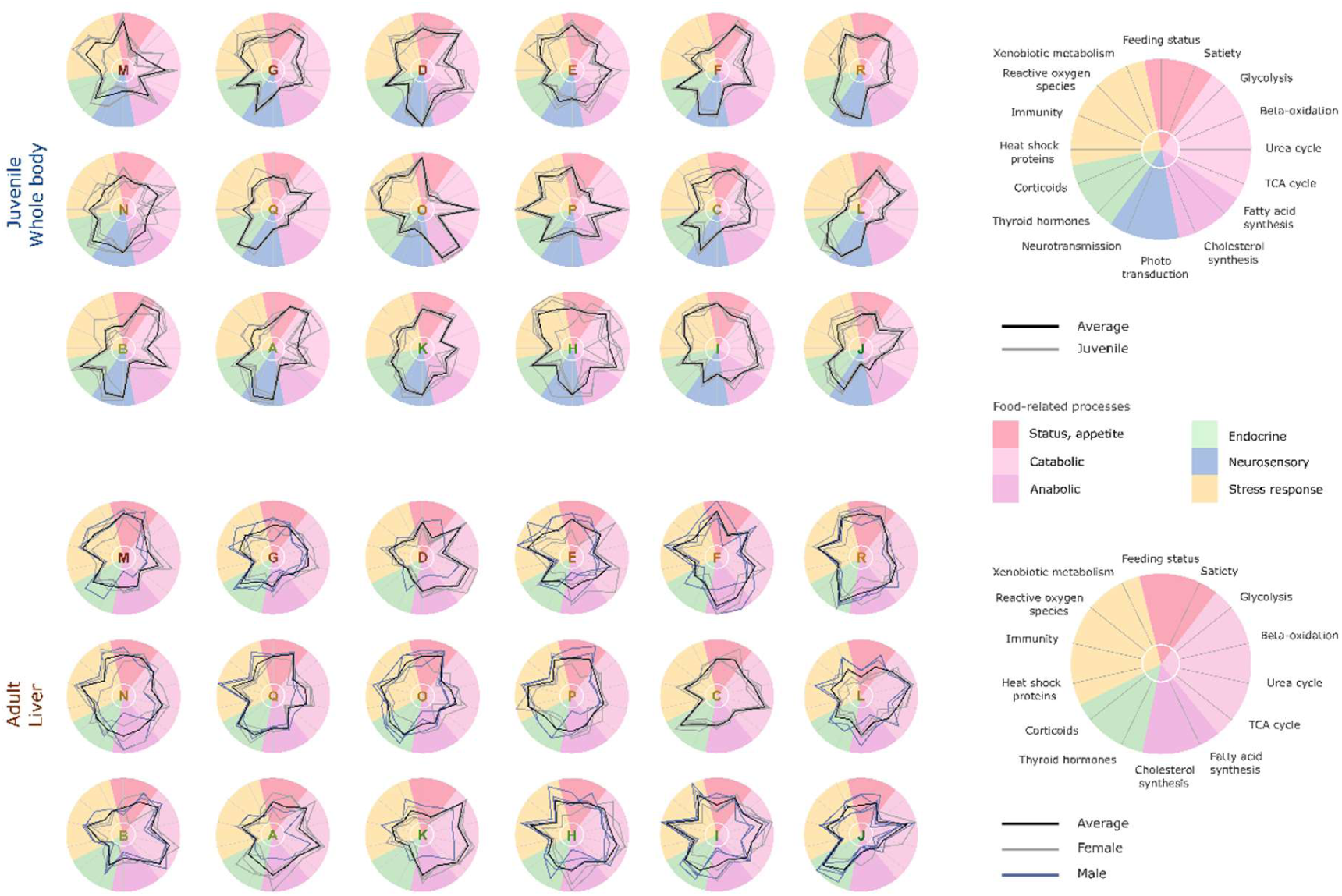
Radar plots presenting the status of juvenile whole-body (top) and adult liver (bottom) across key biological processes. The coordinates were calculated through an orthogonal projection of each sample onto the main axis of site-wise variance based on the top dimensions representing at least 50% of the variance of PCA performed for the following subsets of genes: satiety, glycolysis, beta-oxidation, urea cycle, TCA cycle, fatty acid synthesis, cholesterol synthesis, neurotransmission (juvenile only), photoreception (juvenile only), thyroid hormones, corticoids, heat shock proteins, immunity, reactive oxygen species, xenobiotic metabolism. The feeding status of each sample, as derived from the analysis from Fig. 3, is also presented.

Transcriptomes are complex datasets, mixing genes that can be snapshots of stress with those conveying messages about long-term conditions. The comparison of juvenile whole-body and adult liver profiles allowed us to discern timescales at which environmental factors affect gene expression. For instance, the two most urban sites, G and M, had similar radars for adult livers but not for juvenile whole-body samples, with differences in corticoids, neurotransmission, and energy metabolism (also see the respective PCAs on Fig. 2b,c). We hypothesise that juveniles from G benefited from its more open reef morphology with frequent water renewal, that can wash away waterborne stressors. In contrast, adults that have been exposed to local conditions for longer durations could have integrated chronic stress, particularly in terms of immune response and neurotransmission. Landscape transcriptomes can thus yield time-integrated differences across environments that may be deemed similar, thereby making the ‘invisible’ visible.

### Divergent strategies detected in organisms across environments

Grouping multiple biological processes in the same visualisation enabled the detection of distinct strategies depending on developmental stage, environmental conditions, and feeding. For instance, well-fed juveniles suppressed satiety-signalling genes; this can boost food intake and thus growth (Extended Data Fig. 6). Similarly, energy metabolism shifted from catabolic to anabolic processes with increasing feeding status: juveniles and adults with low-food status (*e.g.,* juveniles from A, B, F, L; adults from A, B, C, H, I, J; Fig. 3) privileged glycolysis, which allows for efficient energy conversion from sugars; in some sites, β-oxidation, which can use fatty reserves to maintain energy homeostasis, was also upregulated (juveniles from B, L; adults from C, H, I, J). Inversely, well-fed fish showed high fatty acid and cholesterol synthesis (highest in O in juveniles, also in D, E, H, I, K, M, N, P; for adults, in F, N, R). A switch in energy investment strategy was detected in high-stress environments: well-fed adult fish showed low metabolism-related gene expression in a trade-off for high immune response in M, and with thermal stress in O and P. We therefore show that landscape transcriptome data provide a rich framework for testing the ecological relevance of physiological regulations identified in experimental animal models.

### Typical pollution marker genes show limited reliability in Okinawa’s *C. cyanea*

As a last step in our elucidation of environmental and human impacts on fish transcriptomes, we assessed common marker genes of pollution, including immunity, endocrine processes, xenobiotic metabolism (involved in the removal of chemical pollutants), and neurotransmission. While we found a close association between immune response genes and human presence (Fig. 2e, 4), most marker genes responded to other drivers.

Starting with endocrine processes, we identified an upregulation of nuclear thyroid hormone receptors under high temperature, overriding any other environmental driver; this is coherent with previous data from coral reef fish^29^. Other thyroid and corticoid-related genes also responded mostly to temperature, although, for juveniles, *dio3b* – which acts by deactivating thyroid hormones in target tissues – and genes in the neuroendocrine cascade leading to cortisol release, notably corticotropin-releasing hormone receptors, pro-opiomelanocortin B, and ferredoxin, were upregulated under low food conditions (Extended Data Figs. 7, 8). Interestingly, *hsd11b1*, a gene involved in the conversion of the inactive cortisone to the active cortisol, was upregulated in juveniles from the natural sites H and I. This indicates an acute stress response, potentially linked to the strain of capture on the fish, which may be dampened elsewhere due to chronic stress.

We found feeding status, rather than pollution, to be linked to xenobiotic metabolism at both stages (Fig. 4; Extended Data Fig. 9). This could be because many of these genes, involved in the modification, conjugation, and excretion of compounds, have a pleiotropic role and intervene in endogenous processes dependent on food-derived energy. Lastly, neurotransmission and immune defence mechanisms, which are energetically costly^30,31^, were suppressed under low food conditions in juveniles (Extended Data Fig. 9).

The tolerance of *C. cyanea* to a large range of conditions and its ubiquitous presence along Okinawa’s coastline allowed us to extract environmental signals in juvenile whole-body and adult liver transcriptomes, revealing intricate patterns of gene expression depending on local conditions, developmental stage, tissue, sex and size. We identified which genes and biological processes respond to human activities, and importantly, which do not. While thermal stress and food availability could be confounding factors, the absence of clear signal from ‘classic’ pollution marker genes may also be due to the choice of species: *C. cyanea* are omnivorous and planktivorous fish near the base of the trophic food web, and whilst territorial and living in shallow waters, they are not benthic fish. They might therefore not bioaccumulate toxicants to levels leading to a significant transcriptomic signature of chemical stress in typical marker genes, notably those involved in xenobiotic metabolism. Comparing the signals obtained with ubiquitous species higher in the food chain or with benthic feeders could be of interest.

## Discussion and outlook

Our study of the localised gene expression response of a common fish species in Okinawa is a step towards building a generalisable framework for environmental state awareness, or ‘environmental phenotyping’, using transcriptomes, with implications for global ecosystem monitoring. In the case of *C. cyanea*, further studies will be required to further elucidate temporal and seasonal variability. In addition, the analysis of multiple tissues at different developmental stages would provide insight into more biological processes as well as help to identify carry-over effects throughout development. Future studies may benefit from detailed environmental characterisations, including long-term water quality data series and measurements of chemical pollution. Lastly, generating additional laboratory-based transcriptomes (*e.g.,* temperature, salinity, chemical compounds) could be an efficient method to isolate the footprint of different stressors in the field.

Landscape transcriptomics provided an unparalleled depth of information about the status of the blue damselfish *C. cyanea* on Okinawa’s coastal zones. Using semi-randomised sampling and complementary laboratory experiments, we identified the combined effects of environmental parameters and the human footprint on transcriptomes and found biological trade-offs in the choice of a habitat. With sufficient genomic resources and environmental information for baseline analyses, our methodology can be generalised to sentinel species across any ecosystem and holds huge potential for environmental monitoring.

## Materials and Methods

### Study location

The subtropical island of Okinawa, Japan, is the largest island of the Ryukyu archipelago. It has an area of approximately 1,200 km² and a population of 1.38 million ^1^. Most inhabitants live in the southern half of the island. The average ocean temperature is 25°C through the action of the warm Kuroshio current, and the island is surrounded by a fringing and patch reef system, which has a length of approximately 380 km. Coral communities represent an area of 69.8 km^2^ ^1^.

Okinawa is an ideal location for this study, with coastal ecosystems heavily influenced by nearby human activities and a range of contrasted coastal ecosystems along different zones, from dense urban areas near the city of Naha in the south, to the Yanbaru forest in the north. The water quality in Okinawa is affected by human activities. Rivers in urban zones were found to have high ammonium and chemical compound concentrations, notably of pharmaceuticals, with compounds such as bezafibrate, caffeine, clarithromycin, and sulfamethoxazole found at higher levels than the 90th percentile across 109 major Japanese rivers ^2^. Endocrine disruptors such as nonylphenol, Bisphenol A, and per- and polyfluoroalkyl substances have been measured in rivers and reefs with a relationship with the density of human activity ^3,4^. Agricultural zones of Okinawa are linked with nitrite, nitrate, ammonium, and red soil runoff^5^. Human presence is also associated with habitat degradation and coastal hardening to protect shores from erosion and reclaim fringing reefs ^6,7^.

### Environmental data collection

Eighteen sites were visited from May to July 2023 along the coastline of Okinawa (Supplementary Table 1,2), during the reproductive season, which is when juvenile *C. cyanea* can be collected. On each site, using a multiparameter probe (RINKO-Profiler CTD logger), water temperature, salinity, dissolved oxygen, turbidity, and chlorophyll-α were measured.

Three replicate 25m long x 4m wide transects were performed to record the fish community. These transects were positioned parallel to the shore, ten meters apart, in at least one meter of water. Transects could not be conducted at two sites, Uka (J) and Kayo (K), due to limited water depth or strong currents. Two passes were performed per transect; mobile, more visible fishes were recorded during the first pass and more cryptic fishes during the second pass ^8^. All fish were identified to the species level. The average fish density (number of fish per m²) and species richness (number of species per m²) were calculated. Additionally, the substrate cover on each site was assessed across the three transects using a point-intercept method, with the type of substrate (live coral, dead coral, macroalgae, seagrass, rubble, hard substrate, sand) recorded every meter.

Land use in the watersheds near each site was assessed based on a publicly available dataset released in January 2023 ^9^. The percentage of urban, agricultural, and natural zones within a radius of 1km of the sites (excluding water) was calculated for each site.

Environmental parameters (salinity, temperature, dissolved oxygen, turbidity, chlorophyll-alpha, substrate cover, fish density, number of species, and Shannon diversity index) were normalised across sites using *scale* of the *base* package in R for all analyses.

The full dataset is provided in the Supplementary Information.

### Study species

The blue damselfish *Chrysiptera cyanea* is a coral reef damselfish characterised by its bright blue pigmentation (Fig. 1A). It occurs on shallow (<10 meters) subtropical and tropical reefs of the Indo-Pacific and can most generally be found around rubble and corals in subtidal reef flats and tide pools (Fig. 1B). It is one of the most abundant fish along the coastline of the island of Okinawa, Japan, and can be found from urban river mouths to isolated beaches ^10^. *C. cyanea* is territorial and lives in small to large schools that typically consist of a few males and several females ^11,12^. The species displays protogynous hermaphroditism, with females turning into males. In Okinawa, during the reproduction season (typically May-August; *49*, *50*), large numbers of juveniles can be found in clusters with or without adults.

### Fish collection and sample preparation

On 18 sites along the coast of Okinawa, 5 juvenile and 5 adult *C. cyanea* (4 on site B) were collected using hand nets and snorkelling or SCUBA diving. In total, 90 juveniles, 49 females, and 40 males were collected. The fish were euthanised in a 200mg/L Tricaine Methanesulfonate (MS222) solution following the guidelines for animal use issued by the Animal Resources Section of OIST Graduate University. Juvenile fish were photographed and directly placed in RNAlater for tissue preservation. They could not be weighed due to field logistics. The average size of the juveniles was 9.6 ± 0.7 mm SL [range of 7.9-1.2 mm SL]. There was no significance difference in fish standard length across sites. Adult fish were photographed, weighed, and dissected on-site. The livers were preserved using RNAlater. The average size of the females was 34.5 ± 5.3 mm SL and of males was 43.8 ± 4.6 mm SL [23.5 – 56.0 mm SL]. The Fulton’s body condition factor (total length in cm divided by cube of the weight in grams) of the females was 1.73 ± 0.31 and of the males was 1.66 ± 0.23 [1.14 – 2.60]. There was no significance difference in Fulton’s body condition factor across sites for females and males. The characteristics of each fish from the field are listed in Supplementary Tables 3 and 4.

### Feeding experiment

The effect of food availability on the transcriptomic profile of *C. cyanea* was assessed with two experiments, for juveniles and adults. The experiment lasted 3 days in juveniles and 14 days in adults due to the difference in tolerance to fasting at the distinct developmental stages.

For juvenile fish, juveniles collected in Apogama, Okinawa, Japan (26.50003 N, 127.84261 E), on July 30^th^, 2024 were kept at the OIST Marine Science Station. After one night of acclimation, they were placed in two tanks with 14 juvenile fish per tank. Water input came from pumped seawater. Two treatments were used: fasted and fed. In one tank, the fish were fed normal amounts of food for three days (dry pellets twice per day, ASIN number: B0BFJ9MY8P); in the other tank, the fish were not fed and a mesh filter was used on the water inlet to remove particles and plankton. After three days, each fish was weighed, measured, euthanised with MS222, and dissected. The whole body of each individual was preserved in RNAlater. Of these samples, seven per treatment (14 in total) were used for RNA sequencing (Supplementary Table 5). The average weight of the sequenced fish at the end of the experiment was 33 ± 9 mg for the fasted group and 77 ± 22 mg for the fed group. Fulton’s condition index was 1.36 ± 0.20 for the fasted group and 2.00 ± 0.21 for the fed group.

For adult fish, female *C. cyanea* collected in Sesoko, Okinawa, Japan (26.65056 N, 127.8549 E), on September 30^th^, 2022 were kept at the OIST Marine Science Station. After an acclimation period, on November 23^rd^, 2022, they were placed in two tanks with eight fish per tank. Water input came from pumped seawater. The experiment had two groups of fish: fasted and fed (control). In one tank, the fish were fed normal amounts of food for two weeks (dry pellets twice per day, ASIN number: B0BFJ9MY8P); in the other tank, the fish were not fed for two weeks. The average weight of the fish at the start of the experiment was 2.3 ± 0.7 g for the fasted group and 2.3 ± 0.9 g for the fed group. After one week, each fish was weighed. The average weight of the fish after one week was 2.1 ± 0.8 g for the fasted group and 2.4 ± 1.1 g for the fed group. After two weeks, on December 6^th^, 2022, each fish was weighed, measured, euthanised with MS222, and dissected. The livers were preserved in RNAlater. Of these samples, seven per treatment (14 in total) were used for RNA sequencing (Supplementary Table 6). The average weight of the sequenced fish after two weeks, at the end of the experiment, was 2.0 ± 0.7 g for the fasted group and 2.4 ± 0.8 g for the fed group. Fulton’s condition index was 1.70 ± 0.09 for the fasted group and 2.04 ± 0.22 for the fed group. The Hepatosomatic index (100 x liver weight over the total weight of the fish) was 0.80 ± 0.14 for the fasted group and 1.55 ± 0.27 for the fed group.

### RNA extraction, library preparation, sequencing, and data processing

RNA extractions were performed using the Maxwell® RSC simplyRNA tissue kits and an automated Maxwell® RSC instrument based on the manufacturer’s recommendations (Promega, Cat. No. AS2000, Wisconsin, USA). The quality and concentration of RNA were assessed with an Invitrogen Qubit Flex benchtop fluorometer and an Agilent 4200 TapeStation. Library preparation was performed using the NEBNext Ultra II Directional RNA library prep kit for Illumina (New England BioLabs, USA). Sequencing was done on an Illumina NovaSeq6000 platform at the OIST Sequencing section. Paired end reads (2 x 150 bp) were obtained.

Quality control of the RNA sequencing data was performed with FastQC ^15^. Following this quality check, the data was processed by trimming the adapters used by Illumina on each transcript as well as dropping low quality sections with Trimmomatic 0.39 ^16^. The sequences were aligned to the reference transcriptome for this species ^17^ and the number of transcripts per gene were quantified using Kallisto 0.46.2 ^18^. The percentage of reads mapping against the reference transcriptome was 68.1 ± 3.8 (mean ± sd) for field juveniles, 81.7 ± 7.1 for field livers, 58.5 ± 3.5 for feeding experiment juveniles, and 73.3 ± 1.7 for feeding experiment livers.

### RNAseq data analysis

All analyses were performed on the transcriptomes of juvenile whole-body and adult liver samples separately. All data analysis and figure generation scripts are available on FigShare. R 4.3.3 was used. Plots were generated using *ggplot2* 3.5.1*, ggtern* 3.5.0*, ggrepel* 0.9.5*, ggradar* 0.2*, pheatmap* 1.0.12 in R. A threshold of p < 0.01 was used throughout the study unless otherwise specified. Principal Component Analyses (PCA) were obtained with the *prcomp* function from *stats* 4.3.3. Statistical tests included the non-parametric Kruskal-Wallis Rank Sum test and linear mixed effect models with environmental variables, fish size or sex as fixed effects and site as a random effect, using the top parameters identified with the *regsubset* model selection function of *leaps* 3.2.

Using all field samples as input, gene counts were normalised using *DESeq2* 1.36.0 ^19^. Genes with more than 10 counts in at least two sites were kept. Following variance stabilising transformation (vsd) with *DESeq2,* the matrix of the gene expression level for each gene in the field samples was used as the basis for the analysis. A second vsd-normalised matrix combining the field and feeding experiment samples together was also used, keeping genes with more than 10 counts in at least two sites or feeding treatments.

Using *DESeq2,* the number of differentially expressed genes in pairwise site comparisons was estimated at the threshold of p < 0.01 and absolute fold change of 1.5 (logFC of 0.58). The number of genes responding to sex in the adult liver samples was also calculated. Using *DRomics* 2.6-2 ^20^, the number of genes responding linearly or quadratically to various environmental factors (percentage of natural zones near each site, temperature, sampling time, salinity, chlorophyll-α) as well as size for juvenile fish was calculated at a significance threshold of p < 0.01.

An annotated genome is available for *Chrysiptera cyanea* with gene names identified for 26,300 genes using BLAST ^17^. Manually curated sets of genes corresponding to key biological processes – appetite regulation, glycolysis, beta-oxidation, urea cycle, TCA cycle, fatty acid synthesis, cholesterol biosynthesis, phototransduction, neurotransmission, thyroid hormones, corticoids, reactive oxygen species – were used ^21^. In addition, genes associated with the immune response, heat shock, and xenobiotic metabolism were identified for the analysis. Module analysis based on subsets of genes was performed with *WGCNA* 1.72-5 ^22^. GO terms (*org.Dr.eg.db*) were analysed semantically using *GoSemSim* 2.28.1 ^23^. KEGG terms were annotated using BlastKOALA ^32^.

### Meta-transcriptomic analysis

To estimate diet in juvenile *C. cyanea*, using Kaiju 1.10.0 ^24^, juvenile whole-body reads that did not map to the reference *C. cyanea* transcriptome were compared with 2,462 reference NCBI protein sequences available for the following taxa: Artemia (NCBI ID: 6660), Chlorophyta (3041), Ctenophora (10197), Cyanobacteria (1117), Dinoflagellata (2864), Embrophyta (3193), Fungi (4751), Haptophyta (2830), Ochrophyta (2696291), Pancrustacea (197562), Rhodophyta (2763), Tunicata (7712). The proportion of reads in each sample corresponding to different taxonomic families was identified.

### Feeding and field experiment analysis

To compare the feeding and field samples, a PCA was performed on the feeding experiment samples only (*i.e.,* fish that were fasted or fed over 3 days for the juvenile experiment, and 14 days for the adult experiment). After identifying the top 10% of genes with the highest loading along PC1, a second PCA was generated based on the subset of genes for the feeding experiment and field samples together. As the field samples fell between the feeding samples along PC1, the PC1 coordinate of each sample was obtained as an indicator of each sample’s feeding status. This yielded five datapoints per site and sample type (four for adult livers from site B).

### Radar plots

Radar plots were produced to summarise the dimensionality in vsd-transformed gene expression levels for various subsets of genes as obtained using Principal Component Analysis (PCA). For each PCA, the minimum number of dimensions accounting for at least to 50% of the total variance was calculated. The average coordinates of each site were calculated for these top dimensions, and the main axis of variance along these coordinates was obtained. The orthogonal projection of all samples along this main axis of variance was then calculated, yielding a segment bounded by the two most extreme samples. The gene expression levels of the two sites which had the most extreme average projections along the segment was compared. After determining which site had the most genes showing lower expression levels, the sample at the extremity of the vector closest to this selected site was set to have a value of 0, while the opposite extremity was set to have a value of 1 (sketch of the method provided in Supplementary Materials). For most biological processes screened to obtain the radar plots, there was a linear correlation between the calculated coordinate of each sample and the mean expression level of the genes within the set (Supplementary Materials), although exceptions were noted (*e.g.,* phototransduction in juveniles, where the set of genes showed a clear difference in expression pattern between morning and afternoon sampling on the PCA, but some genes were upregulated while others were downregulated). The relationship between radar plot coordinates and gene expression was also checked with heatmaps (examples in the Supplementary Materials). If expression patterns with various possible environmental drivers were found, the data structure was further dissected with module analysis tools (*e.g.,* for xenobiotic metabolism and environmental stress, Supplementary Materials).

### Population genetics

For juvenile whole-body transcriptomes, in which a higher number of genes was expressed than in the adult liver samples, the raw sequencing reads were mapped to the reference genome using 2-pass mapping using STAR 2.7.9a followed by data cleaning with Picard (AddOrReplaceReadGroups, MarkDuplicates). GATK 4.4.0.0 was then used for variant calling following the pipeline from Genomics Core at NYU CGSB ^25^. SNPs were filtered to exclude low quality variants. Principal Component Analysis was performed using PLINK, removing samples with over 25% of missing genotype data (11 samples out of 90) and SNPs with missing genotype rates higher than 25% (9,040 variants retained). Discriminant Analysis of Principal Components (DAPC) was performed in R using *adegenet 2.1.10*. No significant population structure was found across five zones around the island – northwest, north, east, south, west (Supplementary Materials).

## Acknowledgements

We thank Elise Billoir for data analysis support with DRomics, Manon Mercader and Erina Kawai for fieldwork support, James Hutasoit for experimental support, Brandon Conlon for scientific writing advice, the OIST Sequencing Section for library preparation and sequencing, and the Scientific Computing and Data Analysis section of Core Facilities at OIST. We also thank James Reimer, Manon Mercader, Eugene Myers, and Daniel Rokhsar for fruitful discussions about the project.

## Author contributions

Conceptualization: EG, VL

Methodology: EG, SM, MH, VL

Investigation: EG, SM, ZC, CAS, JZ, HT, YG, MH, VL

Visualization: EG

Funding acquisition: EG, VL

Project administration: SM, VL

Supervision: SM, MH, VL

Writing – original draft: EG

Writing – review & editing: EG, YG, MH, VL

## Data and code availability

The transcriptome sequencing reads are deposited in the NCBI GenBank database under the BioProject IDs: PRJNA1254330 (field samples), PRJNA1271234 (juvenile feeding samples), PRJNA1168266 (adult feeding samples). All files and codes used in this manuscript for data analysis and visualisation are available at Figshare (doi: 10.6084/m9.figshare.27966972).

## Ethics

Ethical approval for the feeding experiment was granted by the Okinawa Institute of Science and Technology (ACUP-2023-014).

## Additional information

Supplementary Information is available for this paper. Correspondence and requests for materials should be addressed to Emma Gairin.

## Supplementary Information

**Table S1:** Site name, coordinates, date and hour of sampling, percentage of urban, agricultural, and natural zones within a kilometre of each site, and environmental parameters (temperature, salinity, chlorophyll-α, turbidity, dissolved oxygen).

**Table S2:** Substrate characteristics (percentage of live coral, dead coral, macroalgae, seagrass, rubble, sand, rock) and fish community description (density of juveniles and adults of all species, density of *Chrysiptera cyanea* juveniles and adults, number of species, and Shannon diversity index).

**Table S3:** Sample description of the juveniles from the field.

**Table S4:** Sample description of the adults from the field.

**Table S5:** Sample description of the juveniles from the feeding experiment.

**Table S6:** Sample description of the adults from the feeding experiment.

**Table S7:** Spearman correlation coefficient between the projected coordinate of each sample along the main axis of variance across the top dimensions of Principal Component Analyses for subsets of genes, and the mean expression level of the genes.

**Extended Data Fig. 1: Drivers of the overall data structure in two Principal Component Analyses (PCA), using all genes with counts above 10 in at least two sites, for juvenile and liver samples separately.** For juveniles (**a**) and adults (**b**), bar charts of the percentage of contribution to the variance of the top 6 PC dimensions, corresponding to over 50% of the total variance. Tile plots presenting the significance of the linear relationship between various environmental parameters and PC coordinates. The tests were linear mixed effect model with site as a random effect and variables selected from preliminary linear models with lowest BIC value using *regsubset* (*leaps* package in R). **c**, Relationship between PC1 (PCA with all genes with counts above 10 in at least two sites for juvenile samples) and the standard length of the juvenile fish. **d**, Principal Component Analysis of the expression of all genes with normalised counts (from DESEQ2) above an average of 10 in at least two sites, for adult liver samples, *i.e.,* for 17,868 genes. Triangles correspond to female livers and circles to male livers. Sex was assigned based on the tail colour of the fish at sampling (females have transparent caudal fins while males have opaque blue caudal fins). The bottom panel is a violin plot indicating the distribution of the male and female samples along PC1. There is a significant difference in PC1 values depending on sex (Kruskal-Wallis rank sum test, χ^2^ = 40.4, df = 1, p < 10^-9^).

**Extended Data Fig. 2: Feeding experiment results for the juvenile and liver samples. a**, Volcano plot displaying the number of significantly differently expressed genes in the whole body (absolute Log2 fold change > 0.58, p < 0.01) from DESEQ2 for the juvenile feeding experiment (no food *vs.* fed). **b**, Volcano plot displaying the number of significantly differently expressed genes in the liver (absolute Log2 fold change > 0.58, p < 0.01) from DESEQ2 for the adult feeding experiment (no food *vs.* fed). **c**, Venn Diagram displaying the number of differentially expressed genes (absolute Log2 fold change > 0.58, adjusted p < 0.01) from DESEQ2 in the juvenile and liver samples, with the genes in common shown based on the intersection of the different diagram cells. **d**, Principal Component Analysis of the gene expression level across all genes (with mean expression above 10 counts) for the juvenile feeding experiment samples (7 juvenile fish with no food, 7 fed for 3 days). **e**, Principal Component Analysis of the gene expression level across all genes (with mean expression above 10 counts) for the liver feeding experiment samples (7 adult fish with no food, 7 fed for 3 days).

**Extended Data Fig. 3: Most common GO terms across the top 10 percent genes driving the loading of dimension 1 on the Principal Component Analysis (PCAs) of the feeding experiment samples for juveniles (a) and adult liver (b).** The PCAs are presented on Fig. 3. GO terms were trimmed based on semantic similarity at a threshold of 0.7 using GOSemSim and the most unique terms are presented here.

**Extended Data Fig. 4: Most common GO terms across the top 10 percent genes driving the loading of dimension 2 on the Principal Component Analysis of the feeding and field samples (Fig. 3b from the main text) in juveniles.** GO terms were trimmed based on semantic similarity at a threshold of 0.7 using GOSemSim and the most unique terms are presented here.

**Extended Data Fig. 5: Sketch of the method to obtain the radar plots presented in Fig. 4**. This is an example where the first two PC dimensions are used. The analysis includes all PC dimensions representing up to 50% of the total variance. **a**, First, the Principal Component Analysis, here based on the normalized expression level of 16 genes linked to corticoids, across 18 sites in Okinawa, with 5 samples per site, is obtained. **b**, The mean PC1/PC2 coordinate of the samples from each of the 18 sites is calculated. **c**, The direction of the main variance in the mean coordinate of the sites is then calculated, here presented as a dashed axis. **d**, The orthogonal projection of all mean site coordinates as well as the projection of each sample onto the main axis of variance is calculated. The two furthermost samples across the main variance axis are set to have values of 0 and 1. Based on this method, the orthogonal projection of all samples and sites was calculated for multiple gene sets. The results are then summarized as a radar plot in the main manuscript.

**Extended Data Fig. 6: Heatmap of the vsd-normalised expression level of 45 appetite-related genes in juvenile field samples.** Rows are clustered based on correlation using the Ward D2 method with *pheatmap*. Environmental (temperature, hour of sampling) and intrinsic fish variables (feeding status, standard length) are indicated above the heatmap. Increasingly dark colours indicate higher values for the radar plot coordinates, feeding status, standard length, and hour of sampling. Temperature is indicated from a blue (low temperature) to red (high temperature) scale.

**Extended Data Fig. 7: Heatmap of the vsd-normalised expression level of 27 thyroid-related genes in juvenile field samples.** Rows are clustered based on correlation using the Ward D2 method with *pheatmap*. Environmental (temperature, hour of sampling) and intrinsic fish variables (feeding status, standard length) are indicated above the heatmap. Increasingly dark colours indicate higher values for the radar plot coordinates, feeding status, standard length, and hour of sampling. Temperature is indicated from a blue (low temperature) to red (high temperature) scale.

**Extended Data Fig. 8: Heatmap of the vsd-normalised expression level of 25 corticoid-related genes in juvenile field samples**. Rows are clustered based on correlation using the Ward D2 method with *pheatmap*. Environmental (temperature, hour of sampling) and intrinsic fish variables (feeding status, standard length) are indicated above the heatmap. Increasingly dark colours indicate higher values for the radar plot coordinates, feeding status, standard length, and hour of sampling. Temperature is indicated from a blue (low temperature) to red (high temperature) scale.

**Extended Data Fig. 9: Results from a module analysis for the juvenile whole-body and adult liver gene expression.** Analysis performed using WGCNA (Weighted Correlation Network Analysis) with a minimum module size of 20 genes for juveniles and 13 genes for adult liver samples (corresponding to the lowest number of genes within a category) and a soft-thresholding power of 9 for juveniles and 6 for livers. 13 modules of genes with similar expression patterns across the samples were retrieved for the juveniles and 8 for the livers. The following categories of genes were used, with 974 genes retrieved in total: heat shock proteins, reactive oxygen species, corticoids, thyroid hormones, neurotransmission, immunity, and xenobiotic metabolism. Environmental and intrinsic fish variables (percentage of natural zones within a kilometre of the site, feeding status as derived from Fig. 3 of the main text, standard length for juveniles, sex for livers, temperature, and hour of sampling) were used in a linear-mixed effect model (with site as a random effect and a selection of fixed effects based on model dredging) to determine the relationship between the gene expression patterns in each module and environmental or intrinsic fish drivers. The resulting t-values are presented. Higher t-values (in red) indicate positive relationships between the gene expression level within the module and the given variable, and inversely (in blue). Top: Percentage of genes of each category retrieved within a given module. Bottom: t-value of a linear mixed-effect model retrieving significant relationships between environmental or intrinsic fish characteristics and gene expression patterns within the modules.

**Extended Data Fig. 10: Environmental characterisation of the sites. a**, Ternary plot presenting the percentage of natural, agricultural, and urban zones within 1km of the sites (excluding water). **b**, Substrate cover on three 25m transects performed parallel to shore near the sampling (maximum depth: 2m). **c**, NMDS analysis of the juvenile fish community structure and contributing environmental variables based on *vegan* function envfit (p < 0.1). **d**, NMDS analysis of the adult fish community structure and contributing environmental variables based on *vegan* function envfit (p < 0.1). No fish community structure was recorded on sites J and K due to logistical constraints.

**Extended Data Fig. 11: First two discriminant functions of a Discriminant Analysis of Principal Components (DAPC) of *C. cyanea* juveniles.** Colours indicate 5 zones: northwest, north, east, south, west. The analysis is based on filtered SNPs (mind = 0.25, geno = 0.25) following variant calling using the Genome Analysis Toolkit (GATK) pipeline.

## References

1. Gissi, E. et al. A review of the combined effects of climate change and other local human stressors on the marine environment. Science of The Total Environment 755, 142564 (2021).

2. Pirotta, E. et al. Understanding the combined effects of multiple stressors: A new perspective on a longstanding challenge. Science of The Total Environment 821, 153322 (2022).

3. Keagy, J. et al. Landscape transcriptomics as a tool for addressing global change effects across diverse species. Molecular Ecology Resources 0, 1–16 (2023).

4. Forio, M. A. E. & Goethals, P. L. M. An Integrated Approach of Multi-Community Monitoring and Assessment of Aquatic Ecosystems to Support Sustainable Development. Sustainability 12, 5603 (2020).

5. Karydis, M. & Kitsiou, D. Marine water quality monitoring: A review. Marine Pollution Bulletin 77, 23–36 (2013).

6. Martinez-Haro, M. et al. A review on the ecological quality status assessment in aquatic systems using community based indicators and ecotoxicological tools: what might be the added value of their combination? Ecological Indicators 48, 8–16 (2015).

7. Rillig, M. C. et al. The role of multiple global change factors in driving soil functions and microbial biodiversity. Science 366, 886–890 (2019).

8. Walker, C. H., Sibly, R. M., Sibly, R. M. & Peakall, D. B. Principles of Ecotoxicology. (CRC Press, Boca Raton, 2016). doi:10.1201/b11767.

9. Martyniuk, C. J. Are we closer to the vision? A proposed framework for incorporating omics into environmental assessments. Environmental Toxicology and Pharmacology 59, 87–93 (2018).

10. Evans, T. G. Considerations for the use of transcriptomics in identifying the ‘genes that matter’ for environmental adaptation. Journal of Experimental Biology 218, 1925–1935 (2015).

11. Johnson, K. J. et al. A Transformative Vision for an Omics-Based Regulatory Chemical Testing Paradigm. Toxicological Sciences 190, 127–132 (2022).

12. Page, T. M. & Lawley, J. W. The Next Generation Is Here: A Review of Transcriptomic Approaches in Marine Ecology. Front. Mar. Sci. 9, (2022).

13. Snape, J. R., Maund, S. J., Pickford, D. B. & Hutchinson, T. H. Ecotoxicogenomics: the challenge of integrating genomics into aquatic and terrestrial ecotoxicology. Aquatic Toxicology 67, 143–154 (2004).

14. Herrera, M. et al. From Genes to Pathways: A Curated Gene Approach to Accurate Pathway Reconstruction in Teleost Fish Transcriptomics. Journal of Experimental Zoology Part B: Molecular and Developmental Evolution **n/a**, (2025).

15. Lecchini, D., Adjeroud, M., Pratchett, M. S., Cadoret, L. & Galzin, R. Spatial structure of coral reef fish communities in the Ryukyu Islands, southern Japan. Oceanologica Acta 26, 537–547 (2003).

16. Komyakova, V. & Swearer, S. E. Contrasting patterns in habitat selection and recruitment of temperate reef fishes among natural and artificial reefs. Marine Environmental Research 143, 71–81 (2019).

17. Pratchett, M. S., Coker, D. J., Jones, G. P. & Munday, P. L. Specialization in habitat use by coral reef damselfishes and their susceptibility to habitat loss. Ecology and Evolution 2, 2168–2180 (2012).

18. Evans, T. G. & Hofmann, G. E. Defining the limits of physiological plasticity: how gene expression can assess and predict the consequences of ocean change. Philosophical Transactions of the Royal Society B: Biological Sciences 367, 1733–1745 (2012).

19. Halpern, B. S. et al. Recent pace of change in human impact on the world’s ocean. Sci Rep 9, 11609 (2019).

20. Halpern, B. S. et al. A Global Map of Human Impact on Marine Ecosystems. Science 319, 948–952 (2008).

21. Japan Aerospace Exploration Agency (JAXA). High-Resolution Land Use and Land Cover Map of Okinawa Island. (2023).

22. Japan Statistics Bureau. POPULATION AND HOUSEHOLDS OF JAPAN 2020. https://www.stat.go.jp/english/data/kokusei/2020/summary.html (2020).

23. Katzir, I., Cokol, M., Aldridge, B. B. & Alon, U. Prediction of ultra-high-order antibiotic combinations based on pairwise interactions. PLOS Computational Biology 15, e1006774 (2019).

24. Lundstedt, T. et al. Experimental design and optimization. Chemometrics and Intelligent Laboratory Systems 42, 3–40 (1998).

25. Gairin, E. et al. The genome of the sapphire damselfish Chrysiptera cyanea: a new resource to support further investigation of the evolution of Pomacentrids. Gigabyte 2024, 0–0 (2024).

26. Oomen, R. A. & Hutchings, J. A. Transcriptomic responses to environmental change in fishes: Insights from RNA sequencing. FACETS 2, 610–641 (2017).

27. Watson, H., Videvall, E., Andersson, M. N. & Isaksson, C. Transcriptome analysis of a wild bird reveals physiological responses to the urban environment. Sci Rep 7, 44180 (2017).

28. Damiani, G., Sebastiano, M., Dell’Omo, G. & Costantini, D. Blood transcriptome analysis of common kestrel nestlings living in urban and non-urban environments. Science of The Total Environment 928, 172585 (2024).

29. Besson, M. et al. Anthropogenic stressors impact fish sensory development and survival via thyroid disruption. Nat Commun 11, 3614 (2020).

30. Bajgar, A. et al. Extracellular Adenosine Mediates a Systemic Metabolic Switch during Immune Response. PLOS Biology 13, e1002135 (2015).

31. Adamo, S. The Integrated Defense System: Optimizing Defense against Predators, Pathogens, and Poisons. Integr Comp Biol 62, 1536–1546 (2022).

32. Kanehisa, M., Sato, Y. & Morishima, K. BlastKOALA and GhostKOALA: KEGG Tools for Functional Characterization of Genome and Metagenome Sequences. J Mol Biol 428, 726–731 (2016).

## Methods references

1. Japanese Ministry of the Environment, Nature Conservation Bureau, Natural Environment Planning Division. Coral Reefs of Japan. 14 (2020).

2. Mano, H. et al. Water quality comparison of secondary effluent and reclaimed water to ambient river water of southern Okinawa Island via biological evaluation. Environ Monit Assess 189, 442 (2017).

3. Camacho, C. G. et al. PFAS surveillance within a highly militarized island: a case study of Okinawa, Japan. Environmental Science: Processes & Impacts 27, 46–51 (2025).

4. Kawahata, H., Ohta, H., Inoue, M. & Suzuki, A. Endocrine disrupter nonylphenol and bisphenol A contamination in Okinawa and Ishigaki Islands, Japan––within coral reefs and adjacent river mouths. Chemosphere 55, 1519–1527 (2004).

5. Shilla, D. J., Mimura, I., Takagi, K. K. & Tsuchiya, M. Preliminary survey of the nutrient discharge characteristics of Okinawa Rivers, and their potential effects on inshore coral reefs. *Galaxea*, Journal of Coral Reef Studies 15, 172–181 (2013).

6. Masucci, G. D. & Reimer, J. D. Expanding walls and shrinking beaches: loss of natural coastline in Okinawa Island, Japan [PeerJ]. PeerJ 7, e7520 (2019).

7. Nakajima, R., Masucci, G. D., Kuba, R., Wee, H. B. & Reimer, J. D. Son of a beach: Coastal development and the loss of natural beaches over time (1919 to 2018) on Okinawa Island, southern Japan. Marine Pollution Bulletin 212, 117459 (2025).

8. Lecchini, D. & Galzin, R. Spatial repartition and ontogenetic shifts in habitat use by coral reef fishes (Moorea, French Polynesia). Marine Biology 147, 47–58 (2005).

9. Japan Aerospace Exploration Agency (JAXA). High-Resolution Land Use and Land Cover Map of Okinawa Island. (2023).

10. Lecchini, D., Adjeroud, M., Pratchett, M. S., Cadoret, L. & Galzin, R. Spatial structure of coral reef fish communities in the Ryukyu Islands, southern Japan. Oceanologica Acta 26, 537–547 (2003).

11. Tamilmani, G. & Gopakumar, G. Chrysiptera cyanea (Quoy & Gaimard, 1825). in 301– 305 (ICAR - Central Marine Fisheries Research Institute, Kochi, 2017).

12. Wacker, S., Ness, M. H., Östlund-Nilsson, S. & Amundsen, T. Social structure affects mating competition in a damselfish. Coral Reefs 36, 1279–1289 (2017).

13. Bapary, J. M. A., Nurul Amin, Md. & Takemura, A. Food availability as a possible determinant for initiation and termination of reproductive activity in the tropical damselfish Chrysiptera cyanea. Marine Biology Research 8, 154–162 (2012).

14. Bapary, M. A. J., Fainuulelei, P. & Takemura, A. Environmental control of gonadal development in the tropical damselfish Chrysiptera cyanea. Marine Biology Research 5, 462–469 (2009).

15. Andrews, S., Biggins, L., Inglesfield, S. & Montgomery, J. Babraham Bioinformatics - FastQC A Quality Control tool for High Throughput Sequence Data. FastQC. A quality control tool for high throughput sequence data. https://www.bioinformatics.babraham.ac.uk/projects/fastqc/ (2019).

16. Bolger, A. M., Lohse, M. & Usadel, B. Trimmomatic: a flexible trimmer for Illumina sequence data. Bioinformatics 30, 2114–2120 (2014).

17. Gairin, E. et al. The genome of the sapphire damselfish Chrysiptera cyanea: a new resource to support further investigation of the evolution of Pomacentrids. Gigabyte 2024, 0–0 (2024).

18. Bray, N. L., Pimentel, H., Melsted, P. & Pachter, L. Near-optimal probabilistic RNA-seq quantification. Nat Biotechnol 34, 525–527 (2016).

19. Love, M. I., Huber, W. & Anders, S. Moderated estimation of fold change and dispersion for RNA-seq data with DESeq2. Genome Biology 15, 550 (2014).

20. Larras, F. et al. DRomics: A Turnkey Tool to Support the Use of the Dose–Response Framework for Omics Data in Ecological Risk Assessment. Environ. Sci. Technol. 52, 14461–14468 (2018).

21. Herrera, M. et al. From Genes to Pathways: A Curated Gene Approach to Accurate Pathway Reconstruction in Teleost Fish Transcriptomics. Journal of Experimental Zoology Part B: Molecular and Developmental Evolution **n/a**, (2025).

22. Langfelder, P. & Horvath, S. WGCNA: an R package for weighted correlation network analysis. BMC Bioinformatics 9, 559 (2008).

23. Yu, G. et al. GOSemSim: an R package for measuring semantic similarity among GO terms and gene products. Bioinformatics 26, 976–978 (2010).

24. Menzel, P., Ng, K. L. & Krogh, A. Fast and sensitive taxonomic classification for metagenomics with Kaiju. Nat Commun 7, 11257 (2016).

25. Khalfan, M. Variant Calling Pipeline using GATK4 – Genomics Core at NYU CGSB. https://gencore.bio.nyu.edu/variant-calling-pipeline-gatk4/ (2020).

