## Supplementary Materials for "Decoding the constraints acting on a coastal fish using landscape transcriptomics"

**Table S1: Site name, coordinates, date and hour of sampling, percentage of urban, agricultural, and natural zones within a kilometre of each site, and environmental parameters (temperature, salinity, chlorophyll-α, turbidity, dissolved oxygen).**

| Site | Site name | Latitude (N) | Longitude (E) | Nearby features | Date | Hour of sampling | Urban (%) | Agricultural (%) | Natural (%) | Temperature (oC) | Salinity (psu) | Chlorophyll-α (µg/L) | Turbidity (FTU) | Dissolved Oxygen (%) |
|---|---|---|---|---|---|---|---|---|---|---|---|---|---|---|
| A | Seragaki | 26.50514 | 127.8801 | Village, forest | 24/05/2023 | 11:00 | 21 | 10 | 69 | 24.2 | 34.5 | 0.3 | 0.7 | 101 |
| B | Kouri | 26.71363 | 128.0197 | Agriculture | 25/05/2023 | 11:00 | 5 | 39 | 56 | 24.7 | 34.4 | 0.2 | 0.8 | 107 |
| C | Nakijin | 26.70474 | 127.9302 | Village, agriculture | 25/05/2023 | 15:00 | 26 | 35 | 38 | 26.0 | 34.0 | 0.5 | 1.2 | 123 |
| D | Toguchi | 26.36946 | 127.7332 | Village, agriculture, military installation | 26/05/2023 | 10:00 | 22 | 70 | 8 | 24.6 | 33.9 | 0.2 | 0.1 | 102 |
| E | Junkyard | 26.33682 | 127.7444 | City, industry, military installation | 26/05/2023 | 14:30 | 42 | 45 | 13 | 25.1 | 34.3 | 0.2 | 0.2 | 103 |
| F | Mizugama | 26.36009 | 127.7398 | City, large river | 29/05/2023 | 12:00 | 61 | 25 | 14 | 25.7 | 34.5 | 0.1 | 0.1 | 120 |
| G | Minatogawa | 26.27478 | 127.7143 | Industrial zone, military installation | 29/05/2023 | 15:00 | 80 | 13 | 7 | 26.9 | 34.4 | 0.2 | 0.1 | 137 |
| H | Adan | 26.80111 | 128.3195 | Forest | 07/06/2023 | 11:00 | 3 | 11 | 86 | 25.8 | 34.5 | 0.1 | 0.0 | 110 |
| I | Oku | 26.85067 | 128.2821 | Forest, pig farm | 07/06/2023 | 15:00 | 1 | 13 | 86 | 27.1 | 34.4 | 0.3 | 0.8 | 148 |
| J | Uka | 26.83113 | 128.2459 | Forest, main road | 07/06/2023 | 17:00 | 5 | 0 | 94 | 28.6 | 30.7 | 0.1 | 0.0 | 162 |
| K | Kayo | 26.54939 | 128.1106 | Forest, village | 09/06/2023 | 10:00 | 3 | 14 | 83 | 25.5 | 34.5 | 0.3 | 1.4 | 101 |
| L | Sesoko | 26.65056 | 127.8549 | Hotel, agriculture | 09/06/2023 | 14:00 | 23 | 39 | 38 | 26.9 | 34.6 | 0.2 | 0.3 | 122 |
| M | Ginowan | 26.27643 | 127.727 | City | 12/06/2023 | 15:00 | 84 | 13 | 2 | 27.8 | 33.4 | 1.1 | 0.7 | 120 |
| N | Oodo | 26.08846 | 127.7102 | Village, agriculture | 11/07/2023 | 11:00 | 11 | 60 | 28 | 28.3 | 34.4 | 0.1 | 0.0 | 106 |
| O | Hanashiro | 26.11425 | 127.7444 | Golf course | 11/07/2023 | 15:00 | 23 | 48 | 29 | 32.0 | 34.4 | 0.4 | 0.6 | 166 |

| | | | | | | | | | | | | | | |
|---|---|---|---|---|---|---|---|---|---|---|---|---|---|---|
| P | Kuba | 26.27971 | 127.8164 | Village, agriculture, power plant | 12/07/2023 | 14:30 | 43 | 21 | 35 | 32.4 | 34.6 | 0.2 | 0.4 | 140 |
| Q | Agarihama | 26.19888 | 127.7708 | City | 12/07/2023 | 16:30 | 55 | 17 | 28 | 30.4 | 34.5 | 2.7 | 36.2 | 125 |
| R | Kin | 26.44667 | 127.9136 | City, military installation | 14/07/2023 | 10:00 | 66 | 14 | 20 | 29.7 | 34.2 | 0.5 | 1.4 | 102 |

**Table S2: Substrate characteristics (percentage of live coral, dead coral, macroalgae, seagrass, rubble, sand, rock) and fish community description (density of juveniles and adults of all species, density of *Chrysiptera cyanea* juveniles and adults, number of species, and Shannon diversity index).**

| Site | Live coral (%) | Dead coral (%) | Macroalgae (%) | Seagrass (%) | Rubble (%) | Sand (%) | Rock (%) | Juvenile density (number of fish per 100 $m^2$) | Adult density (number of fish per 100 $m^2$) | Juvenile C. cyanea density (number of fish per 100 $m^2$) | Adult C. cyanea density (number of fish per 100 $m^2$) | Number of species (juveniles) | Number of species (adults) | Number of species (all stages) | Shannon index (juveniles) | Shannon index (adults) | Shannon index (all stages) |
|---|---|---|---|---|---|---|---|---|---|---|---|---|---|---|---|---|---|
| A | 4 | 4 | 1 | 19 | 45 | 20 | 7 | 272 | 68 | 74 | 13 | 12 | 10 | 14 | 0.8 | 1.3 | 1.0 |
| B | 7 | 4 | 0 | 0 | 23 | 10 | 56 | 85 | 225 | 11 | 39 | 11 | 18 | 19 | 1.7 | 1.6 | 1.8 |
| C | 3 | 4 | 5 | 19 | 32 | 19 | 18 | 42 | 18 | 1 | 0 | 7 | 7 | 9 | 1.0 | 1.3 | 1.6 |
| D | 1 | 0 | 15 | 8 | 27 | 26 | 23 | 186 | 66 | 34 | 7 | 11 | 13 | 15 | 1.3 | 1.7 | 1.7 |
| E | 0 | 0 | 12 | 0 | 47 | 20 | 21 | 319 | 131 | 72 | 26 | 12 | 11 | 15 | 1.0 | 1.2 | 1.3 |
| F | 0 | 0 | 0 | 0 | 16 | 19 | 65 | 544 | 220 | 152 | 21 | 17 | 24 | 28 | 0.8 | 2.0 | 1.4 |
| G | 7 | 1 | 7 | 0 | 19 | 4 | 62 | 182 | 227 | 39 | 1 | 10 | 21 | 22 | 1.0 | 1.9 | 2.1 |
| H | 24 | 13 | 1 | 0 | 16 | 11 | 35 | 201 | 142 | 5 | 10 | 19 | 23 | 29 | 1.5 | 2.4 | 2.3 |
| I | 19 | 9 | 3 | 0 | 15 | 8 | 46 | 506 | 153 | 47 | 3 | 15 | 24 | 30 | 1.1 | 2.5 | 1.9 |
| J | 0 | 0 | 0 | 0 | 0 | 0 | 100 | N/A | N/A | N/A | N/A | N/A | N/A | N/A | N/A | N/A | N/A |
| K | 0 | 0 | 12 | 0 | 0 | 68 | 20 | N/A | N/A | N/A | N/A | N/A | N/A | N/A | N/A | N/A | N/A |
| L | 4 | 1 | 24 | 1 | 24 | 31 | 15 | 168 | 92 | 27 | 2 | 10 | 19 | 20 | 1.3 | 2.4 | 2.2 |
| M | 0 | 0 | 67 | 0 | 16 | 17 | 0 | 164 | 113 | 41 | 13 | 10 | 6 | 9 | 0.8 | 0.7 | 1.1 |

| | | | | | | | | | | | | | | | | | |
|---|---|---|---|---|---|---|---|---|---|---|---|---|---|---|---|---|---|
| N | 3 | 12 | 4 | 0 | 0 | 48 | 33 | 332 | 187 | 52 | 20 | 17 | 25 | 31 | 1.6 | 2.3 | 2.1 |
| O | 15 | 4 | 16 | 0 | 13 | 7 | 45 | 315 | 354 | 71 | 60 | 21 | 26 | 32 | 1.3 | 1.7 | 1.7 |
| P | 0 | 0 | 21 | 4 | 24 | 33 | 18 | 100 | 32 | 1 | 2 | 12 | 7 | 10 | 1.5 | 1.3 | 1.8 |
| Q | 0 | 0 | 0 | 0 | 32 | 20 | 48 | 84 | 78 | 1 | 18 | 6 | 9 | 9 | 1.1 | 1.3 | 1.6 |
| R | 17 | 1 | 11 | 0 | 17 | 39 | 15 | 477 | 371 | 38 | 18 | 34 | 19 | 38 | 2.2 | 1.7 | 2.2 |

**Table S3: Sample description of the juveniles from the field.**

| Sample | Site | Standard length (cm) | Total length (cm) |
|---|---|---|---|
| AJ1 | A | 1.10 | 1.55 |
| AJ2 | A | 1.01 | 1.35 |
| AJ3 | A | 1.03 | 1.37 |
| AJ4 | A | 0.93 | 1.14 |
| AJ5 | A | 0.97 | 1.31 |
| BJ1 | B | 1.06 | 1.44 |
| BJ2 | B | 0.92 | 1.24 |
| BJ3 | B | 0.91 | 1.23 |
| BJ4 | B | 0.97 | 1.30 |
| BJ6 | B | 0.92 | 1.28 |
| CJ1 | C | 0.97 | 1.39 |
| CJ2 | C | 0.96 | 1.39 |
| CJ3 | C | 0.96 | 1.26 |
| CJ4 | C | 1.03 | 1.40 |
| CJ5 | C | 0.88 | 1.23 |
| DJ1 | D | 1.00 | 1.34 |
| DJ2 | D | 0.99 | 1.33 |
| DJ3 | D | 0.91 | 1.19 |
| DJ4 | D | 0.89 | 1.22 |
| DJ5 | D | 0.93 | 1.31 |
| EJ1 | E | 1.10 | 1.43 |
| EJ2 | E | 0.99 | 1.34 |
| EJ3 | E | 1.13 | 1.54 |
| EJ4 | E | 0.97 | 1.37 |
| EJ5 | E | 0.99 | 1.33 |
| FJ1 | F | 1.04 | 1.48 |
| FJ3 | F | 0.98 | 1.35 |
| FJ4 | F | 0.91 | 1.22 |
| FJ5 | F | 0.85 | 1.18 |
| FJ6 | F | 0.80 | 1.14 |
| GJ1 | G | 0.94 | 1.30 |
| GJ3 | G | 0.90 | 1.28 |
| GJ4 | G | 0.87 | 1.17 |
| GJ6 | G | 0.96 | 1.31 |
| GJ7 | G | 0.87 | 1.18 |
| HJ1 | H | 1.03 | 1.44 |
| HJ2 | H | 0.96 | 1.36 |
| HJ3 | H | 1.24 | 1.76 |
| HJ4 | H | 1.07 | 1.50 |
| HJ5 | H | 0.90 | 1.34 |
| IJ1 | I | 0.90 | 1.30 |
| IJ2 | I | 1.00 | 1.34 |

| | | | |
|---|---|---|---|
| IJ3 | I | 0.96 | 1.40 |
| IJ4 | I | 1.00 | 1.41 |
| IJ5 | I | 0.98 | 1.31 |
| JJ1 | J | 0.89 | 1.24 |
| JJ2 | J | 1.03 | 1.41 |
| JJ3 | J | 1.00 | 1.38 |
| JJ5 | J | 0.96 | 1.20 |
| JJ6 | J | 0.94 | 1.35 |
| KJ1 | K | 1.00 | 1.40 |
| KJ2 | K | 0.96 | 1.39 |
| KJ3 | K | 0.92 | 1.29 |
| KJ4 | K | 1.00 | 1.28 |
| KJ5 | K | 1.00 | 1.26 |
| LJ1 | L | 0.86 | 1.18 |
| LJ2 | L | 0.92 | 1.30 |
| LJ3 | L | 0.94 | 1.31 |
| LJ4 | L | 0.90 | 1.29 |
| LJ5 | L | 0.79 | 1.11 |
| MJ2 | M | 1.03 | 1.47 |
| MJ3 | M | 1.10 | 1.60 |
| MJ4 | M | 1.04 | 1.50 |
| MJ5 | M | 0.83 | 1.18 |
| MJ6 | M | 0.85 | 1.22 |
| NJ1 | N | 0.92 | 1.28 |
| NJ2 | N | 0.96 | 1.37 |
| NJ3 | N | 0.98 | 1.39 |
| NJ4 | N | 0.93 | 1.25 |
| NJ5 | N | 0.96 | 1.28 |
| OJ1 | O | 1.18 | 1.78 |
| OJ2 | O | 1.03 | 1.52 |
| OJ3 | O | 0.93 | 1.34 |
| OJ4 | O | 0.90 | 1.38 |
| OJ5 | O | 1.03 | 1.55 |
| PJ1 | P | 1.13 | 1.60 |
| PJ2 | P | 0.97 | 1.35 |
| PJ3 | P | 0.89 | 1.25 |
| PJ4 | P | 0.99 | 1.41 |
| PJ5 | P | 1.12 | 1.58 |
| QJ1 | Q | 0.95 | 1.29 |
| QJ2 | Q | 0.93 | 1.29 |
| QJ3 | Q | 0.88 | 1.21 |
| QJ4 | Q | 0.92 | 1.22 |
| QJ5 | Q | 0.88 | 1.29 |
| RJ1 | R | 0.98 | 1.38 |
| RJ2 | R | 0.92 | 1.26 |
| RJ3 | R | 0.95 | 1.34 |

| | | | |
|---|---|---|---|
| RJ5 | R | 0.90 | 1.24 |
| RJ6 | R | 0.90 | 1.27 |

**Table S4: Sample description of the adults from the field.**

| Sample | Site | Sex | Standard length (cm) | Total length (cm) | Weight (g) | Fulton’s condition index |
|---|---|---|---|---|---|---|
| AL6 | A | Female | 2.96 | 4.02 | 1.3 | 2.00 |
| AL7 | A | Male | 4.69 | 6.76 | 5.0 | 1.62 |
| AL8 | A | Female | 3.29 | 4.44 | 1.1 | 1.14 |
| AL9 | A | Female | 3.25 | 4.32 | 1.0 | 1.24 |
| AL10 | A | Female | 3.16 | 4.18 | 1.1 | 1.37 |
| BL6 | B | Female | 3.16 | 4.16 | 1.0 | 1.39 |
| BL7 | B | Male | 5.04 | 6.5 | 3.6 | 1.31 |
| BL8 | B | Female | 3.20 | 4.20 | 1.1 | 1.35 |
| BL9 | B | Male | 4.41 | 5.84 | 3.0 | 1.51 |
| CL6 | C | Female | 3.62 | 4.78 | 2.1 | 1.92 |
| CL7 | C | Female | 4.01 | 5.25 | 3.0 | 2.07 |
| CL8 | C | Female | 4.13 | 5.24 | 2.5 | 1.74 |
| CL9 | C | Female | 3.35 | 4.65 | 1.8 | 1.79 |
| CL11 | C | Female | 4.77 | 5.92 | 3.3 | 1.59 |
| DL6 | D | Female | 4.18 | 5.62 | 2.6 | 1.46 |
| DL7 | D | Female | 5.11 | 6.91 | 4.8 | 1.45 |
| DL8 | D | Female | 3.66 | 4.89 | 1.4 | 1.20 |
| DL9 | D | Male | 3.34 | 4.42 | 1.5 | 1.74 |
| DL10 | D | Female | 3.07 | 4.23 | 1.2 | 1.59 |
| EL6 | E | Male | 4.93 | 6.52 | 4.4 | 1.59 |
| EL7 | E | Male | 4.04 | 5.42 | 2.8 | 1.76 |
| EL8 | E | Female | 3.11 | 4.44 | 1.5 | 1.71 |
| EL9 | E | Female | 3.49 | 4.61 | 1.8 | 1.84 |
| EL11 | E | Male | 4.38 | 6.08 | 3.6 | 1.60 |
| FL6 | F | Male | 4.90 | 6.79 | 5.0 | 1.60 |
| FL7 | F | Male | 4.93 | 6.37 | 3.9 | 1.51 |
| FL8 | F | Male | 4.77 | 6.66 | 4.1 | 1.39 |
| FL9 | F | Female | 3.78 | 5.03 | 2.4 | 1.89 |
| FL10 | F | Female | 3.89 | 5.30 | 2.5 | 1.68 |
| GL6 | G | Female | 4.21 | 5.59 | 2.6 | 1.49 |
| GL7 | G | Male | 4.11 | 5.53 | 2.9 | 1.71 |
| GL8 | G | Male | 4.00 | 5.32 | 2.2 | 1.46 |
| GL9 | G | Male | 4.41 | 6.28 | 3.8 | 1.53 |
| GL10 | G | Female | 3.05 | 4.07 | 1.5 | 2.22 |
| HL6 | H | Male | 4.36 | 6.00 | 4.7 | 2.18 |
| HL7 | H | Female | 3.36 | 4.62 | 1.9 | 1.93 |
| HL8 | H | Male | 4.15 | 5.60 | 3.1 | 1.77 |
| HL9 | H | Male | 4.75 | 6.35 | 3.9 | 1.52 |

| | | | | | | |
|---|---|---|---|---|---|---|
| HL10 | H | Female | 3.07 | 4.37 | 1.8 | 2.16 |
| IL6 | I | Male | 4.27 | 5.76 | 3.0 | 1.57 |
| IL7 | I | Male | 3.65 | 4.94 | 2.3 | 1.91 |
| IL8 | I | Male | 3.21 | 4.43 | 1.7 | 1.96 |
| IL9 | I | Female | 3.17 | 4.47 | 1.7 | 1.90 |
| IL10 | I | Female | 3.25 | 4.40 | 1.3 | 1.53 |
| JL6 | J | Male | 4.20 | 5.83 | 3.4 | 1.72 |
| JL7 | J | Male | 3.95 | 5.55 | 2.7 | 1.58 |
| JL8 | J | Male | 4.45 | 6.09 | 3.4 | 1.51 |
| JL9 | J | Male | 4.15 | 5.63 | 2.4 | 1.34 |
| JL10 | J | Female | 3.40 | 4.74 | 1.8 | 1.69 |
| KL6 | K | Female | 2.94 | 4.18 | 1.1 | 1.51 |
| KL7 | K | Female | 2.70 | 3.90 | 1.0 | 1.69 |
| KL9 | K | Female | 2.83 | 3.87 | 1.0 | 1.73 |
| KL10 | K | Male | 3.91 | 5.64 | 2.7 | 1.50 |
| KL11 | K | Female | 2.92 | 4.15 | 1.1 | 1.54 |
| LL6 | L | Female | 3.00 | 3.84 | 1.0 | 1.77 |
| LL7 | L | Male | 3.92 | 5.31 | 3.0 | 2.00 |
| LL8 | L | Female | 3.30 | 4.57 | 1.6 | 1.68 |
| LL9 | L | Female | 3.40 | 4.72 | 2.3 | 2.19 |
| LL10 | L | Male | 4.54 | 6.30 | 3.8 | 1.52 |
| ML6 | M | Male | 4.71 | 6.15 | 5.6 | 2.41 |
| ML7 | M | Male | 5.60 | 7.71 | 7.5 | 1.64 |
| ML8 | M | Female | 3.84 | 5.08 | 2.8 | 2.14 |
| ML9 | M | Female | 3.80 | 5.12 | 2.3 | 1.71 |
| ML10 | M | Female | 4.20 | 5.46 | 3.1 | 1.90 |
| NL6 | N | Male | 4.20 | 5.78 | 2.8 | 1.45 |
| NL7 | N | Female | 2.83 | 4.10 | 1.1 | 1.60 |
| NL8 | N | Male | 4.18 | 5.73 | 2.4 | 1.28 |
| NL9 | N | Female | 3.35 | 4.5 | 1.4 | 1.54 |
| NL10 | N | Female | 3.72 | 5.19 | 2.6 | 1.86 |
| OL6 | O | Female | 2.91 | 3.95 | 1.6 | 2.60 |
| OL7 | O | Male | 4.95 | 6.75 | 5.4 | 1.76 |
| OL8 | O | Male | 4.25 | 5.63 | 3.4 | 1.91 |
| OL9 | O | Female | 3.24 | 4.69 | 1.9 | 1.84 |
| OL10 | O | Male | 4.38 | 6.24 | 4.2 | 1.73 |
| PL6 | P | Male | 4.37 | 5.87 | 3.3 | 1.63 |
| PL7 | P | Female | 3.69 | 4.87 | 2.4 | 2.08 |
| PL8 | P | Female | 3.57 | 4.71 | 2.4 | 2.30 |
| PL9 | P | Male | 4.12 | 5.54 | 3.1 | 1.82 |
| PL10 | P | Female | 2.35 | 4.24 | 0.9 | 1.18 |
| QL6 | Q | Female | 4.21 | 5.70 | 2.8 | 1.51 |
| QL7 | Q | Female | 3.50 | 4.8 | 2.2 | 1.99 |
| QL8 | Q | Male | 4.85 | 6.56 | 4.9 | 1.74 |
| QL9 | Q | Male | 4.59 | 6.22 | 3.9 | 1.62 |
| QL10 | Q | Female | 3.40 | 4.75 | 2.1 | 1.96 |

| | | | | | | |
|---|---|---|---|---|---|---|
| RL6 | R | Male | 4.16 | 5.60 | 2.4 | 1.37 |
| RL7 | R | Male | 4.70 | 6.20 | 4.2 | 1.76 |
| RL8 | R | Female | 3.38 | 4.58 | 1.6 | 1.67 |
| RL9 | R | Male | 4.37 | 5.80 | 3.5 | 1.79 |
| RL10 | R | Female | 2.89 | 4.03 | 1.2 | 1.83 |

**Table S5: Sample description of the juveniles from the feeding experiment.**

| Sample | Standard length (cm) | Total length (cm) | Weight (mg) | Fulton's condition index |
|---|---|---|---|---|
| Fasted 1 | 0.95 | 1.22 | 20.7 | 1.14 |
| Fasted 2 | 1.14 | 1.42 | 41.2 | 1.44 |
| Fasted 3 | 0.98 | 1.24 | 20.9 | 1.10 |
| Fasted 4 | 1.15 | 1.47 | 39.0 | 1.23 |
| Fasted 5 | 1.00 | 1.29 | 32.9 | 1.53 |
| Fasted 6 | 1.00 | 1.30 | 33.9 | 1.54 |
| Fasted 7 | 1.11 | 1.43 | 46.0 | 1.57 |
| Fed 1 | 1.25 | 1.57 | 65.9 | 1.70 |
| Fed 2 | 1.41 | 1.78 | 108.8 | 1.93 |
| Fed 3 | 1.07 | 1.40 | 53.3 | 1.94 |
| Fed 4 | 1.37 | 1.72 | 108.4 | 2.13 |
| Fed 5 | 1.17 | 1.48 | 75.1 | 2.32 |
| Fed 6 | 1.18 | 1.46 | 66.8 | 2.15 |
| Fed 7 | 1.18 | 1.51 | 63.0 | 1.83 |

**Table S6: Sample description of the adults from the feeding experiment.**

| Sample | Standard length (cm) | Total length (cm) | Weight (g) | Fulton's condition index | Liver weight (mg) | Hepatosomatic index |
|---|---|---|---|---|---|---|
| Fasted 1 | 4.13 | 4.7 | 1.86 | 1.79 | 11 | 0.59 |
| Fasted 2 | 3.77 | 4.43 | 1.43 | 1.64 | 14 | 0.98 |
| Fasted 3 | 3.3 | 3.99 | 1.15 | 1.81 | 8 | 0.70 |
| Fasted 4 | 4.99 | 5.76 | 3.10 | 1.62 | 22 | 0.71 |
| Fasted 5 | 4.1 | 4.93 | 2.07 | 1.73 | 18 | 0.87 |
| Fasted 6 | 3.9 | 4.89 | 1.84 | 1.57 | 15 | 0.82 |
| Fasted 7 | 4.46 | 5.24 | 2.51 | 1.74 | 23 | 0.92 |
| Fed 1 | 3.86 | 4.66 | 1.86 | 1.84 | 28 | 1.51 |
| Fed 2 | 4.17 | 5.02 | 2.68 | 2.12 | 46 | 1.72 |
| Fed 3 | 3.87 | 4.63 | 1.77 | 1.78 | 30 | 1.70 |
| Fed 4 | 3.89 | 4.63 | 2.12 | 2.14 | 25 | 1.18 |
| Fed 5 | 4.27 | 4.97 | 3.01 | 2.45 | 50 | 1.66 |
| Fed 6 | 3.49 | 4.17 | 1.42 | 1.96 | 27 | 1.90 |
| Fed 7 | 4.96 | 5.77 | 3.85 | 2.00 | 47 | 1.22 |

**Table S7: Spearman correlation coefficient between the projected coordinate of each sample along the main axis of variance across the top dimensions of Principal Component Analyses for subsets of genes, and the mean expression level of the genes.**

| Juveniles | | |
|---|---|---|
| Category | Number of genes | Correlation |
| Satiety | 45 | 0.58 |
| Glycolysis | 63 | 0.46 |
| Beta-oxidation | 15 | 0.96 |
| TCA cycle | 33 | 0.95 |
| Urea cycle | 7 | 0.48 |
| Fatty acid synthesis | 46 | 0.19 |
| Cholesterol biosynthesis | 26 | 1 |
| Corticoids | 25 | 0.77 |
| Thyroid hormones | 27 | 0.83 |
| Neurotransmission | 127 | 0.97 |
| Phototransduction | 45 | 0.45 |
| Heat shock proteins | 36 | 0.95 |
| Immunity | 403 | 0.96 |
| Reactive oxygen species | 20 | 0.99 |
| Xenobiotic metabolism | 219 | 0.9 |

| Liver | | |
|---|---|---|
| Category | Number of genes | Correlation |
| Satiety | 19 | 0.24 |
| Glycolysis | 56 | 0.27 |
| Beta-oxidation | 15 | 0.92 |
| TCA cycle | 31 | 0.86 |
| Urea cycle | 6 | 0.75 |
| Fatty acid synthesis | 39 | 0.97 |
| Cholesterol biosynthesis | 26 | 1 |
| Corticoids | 13 | 0.26 |
| Thyroid hormones | 16 | 0.92 |
| Heat shock proteins | 28 | 0.93 |
| Immunity | 273 | 0.96 |
| Reactive oxygen species | 19 | 0.84 |
| Xenobiotic metabolism | 195 | 0.96 |

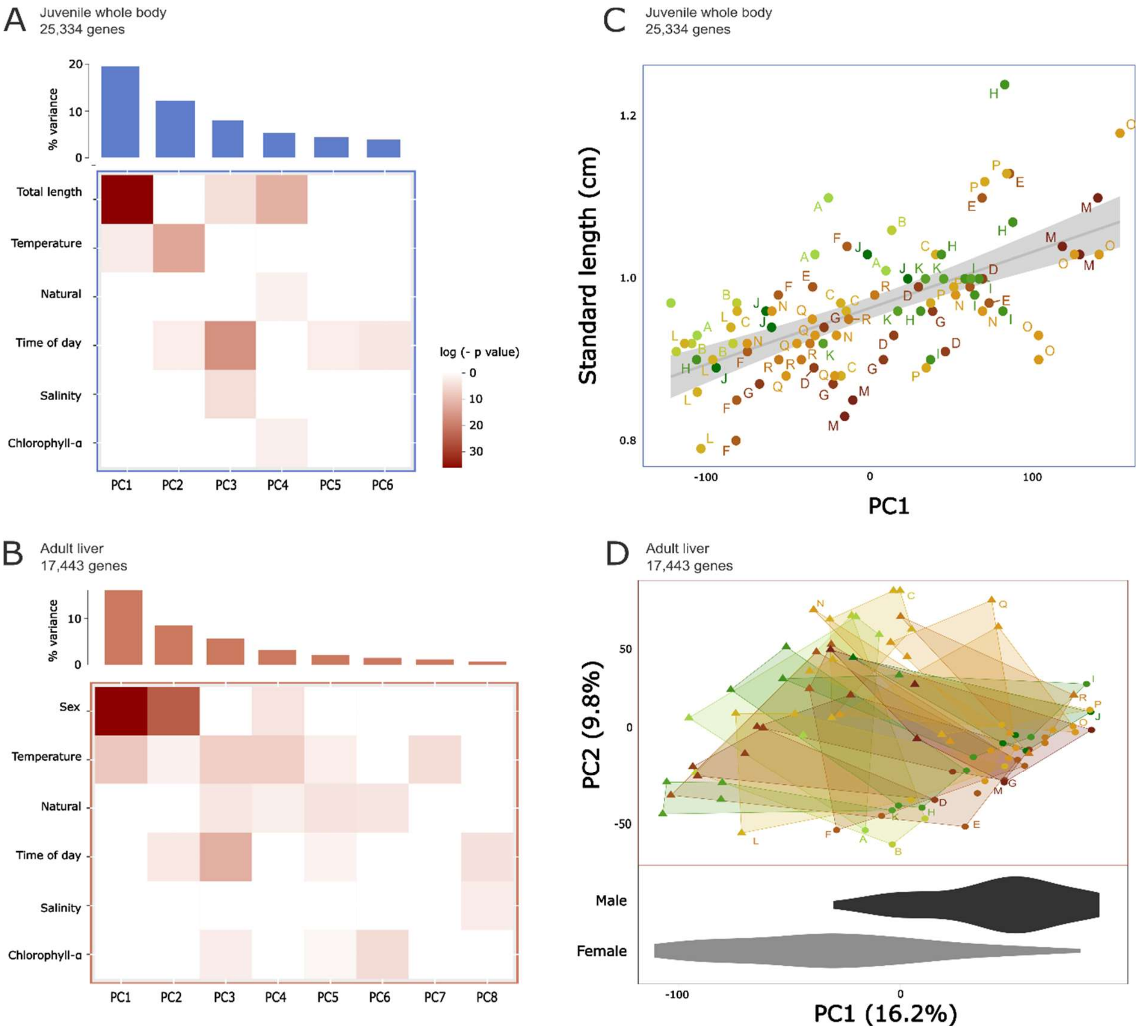


**Extended Data Fig. 1: Drivers of the overall data structure in two Principal Component Analyses (PCA), using all genes with counts above 10 in at least two sites, for juvenile and liver samples separately.** For juveniles (**a**) and adults (**b**), bar charts of the percentage of contribution to the variance of the top 6 PC dimensions, corresponding to over 50% of the total variance. Tile plots presenting the significance of the linear relationship between various environmental parameters and PC coordinates. The tests were linear mixed effect model with site as a random effect and variables selected from preliminary linear models with lowest BIC value using *regsubset* (*leaps* package in R). **c**, Relationship between PC1 (PCA with all genes with counts above 10 in at least two sites for juvenile samples) and the standard length of the juvenile fish. **d**, Principal Component Analysis of the expression of all genes with normalised counts (from DESEQ2) above an average of 10 in at least two sites, for adult liver samples, *i.e.,* for 17,868 genes. Triangles correspond to female livers and circles to male livers. Sex was assigned based on the tail colour of the fish at sampling (females have transparent caudal fins while males have opaque blue caudal fins). The bottom panel is a violin plot indicating the distribution of the male and female samples along PC1. There is a significant difference in PC1 values depending on sex (Kruskal-Wallis rank sum test, $\chi^2 = 40.4$, df = 1, $p < 10^{-9}$).

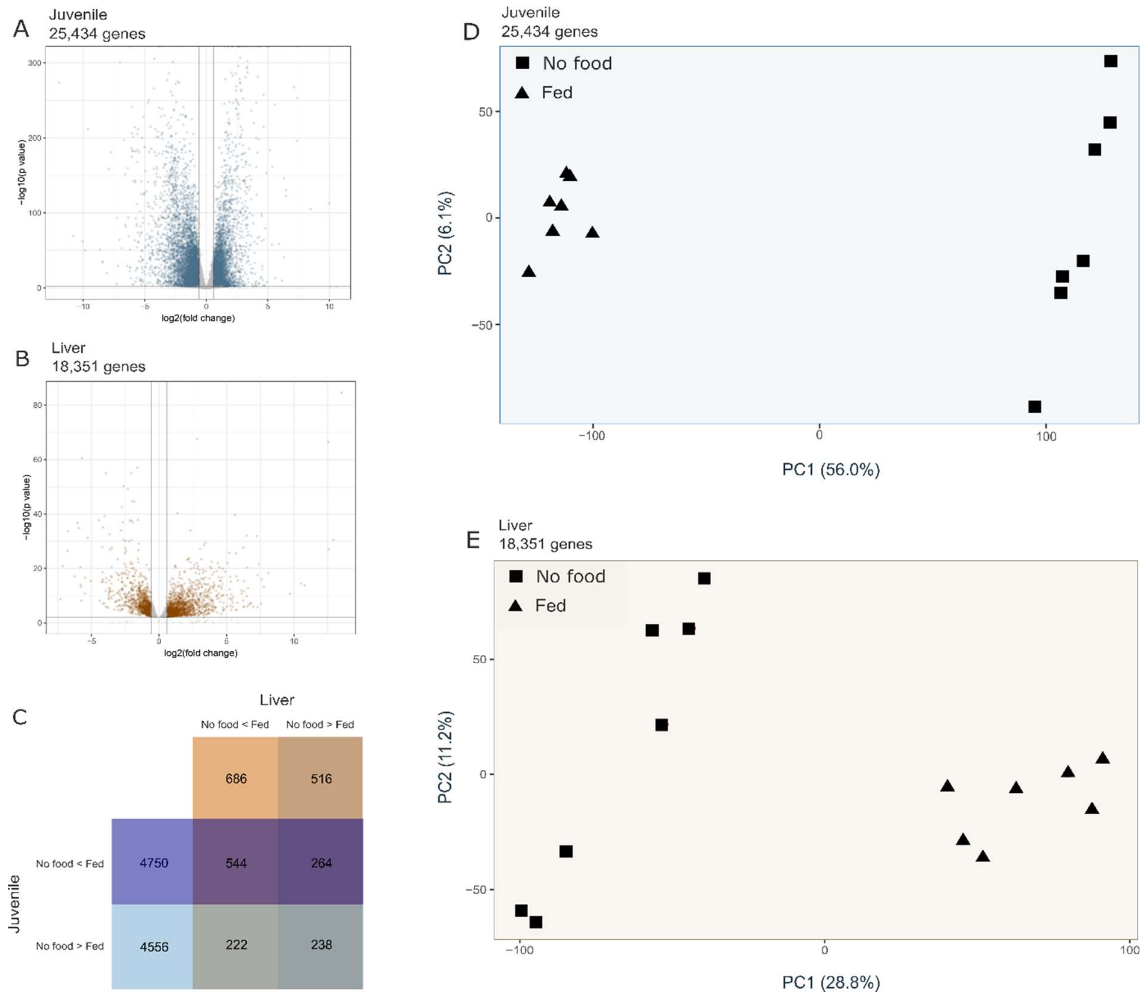


**Extended Data Fig. 2: Feeding experiment results for the juvenile and liver samples. a**, Volcano plot displaying the number of significantly differently expressed genes in the whole body (absolute Log2 fold change > 0.58, $p < 0.01$) from DESEQ2 for the juvenile feeding experiment (no food *vs.* fed). **b**, Volcano plot displaying the number of significantly differently expressed genes in the liver (absolute Log2 fold change > 0.58, $p < 0.01$) from DESEQ2 for the adult feeding experiment (no food *vs.* fed). **c**, Venn Diagram displaying the number of differentially expressed genes (absolute Log2 fold change > 0.58, adjusted $p < 0.01$) from DESEQ2 in the juvenile and liver samples, with the genes in common shown based on the intersection of the different diagram cells. **d**, Principal Component Analysis of the gene expression level across all genes (with mean expression above 10 counts) for the juvenile feeding experiment samples (7 juvenile fish with no food, 7 fed for 3 days). **e**, Principal Component Analysis of the gene expression level across all genes (with mean expression above 10 counts) for the liver feeding experiment samples (7 adult fish with no food, 7 fed for 3 days).

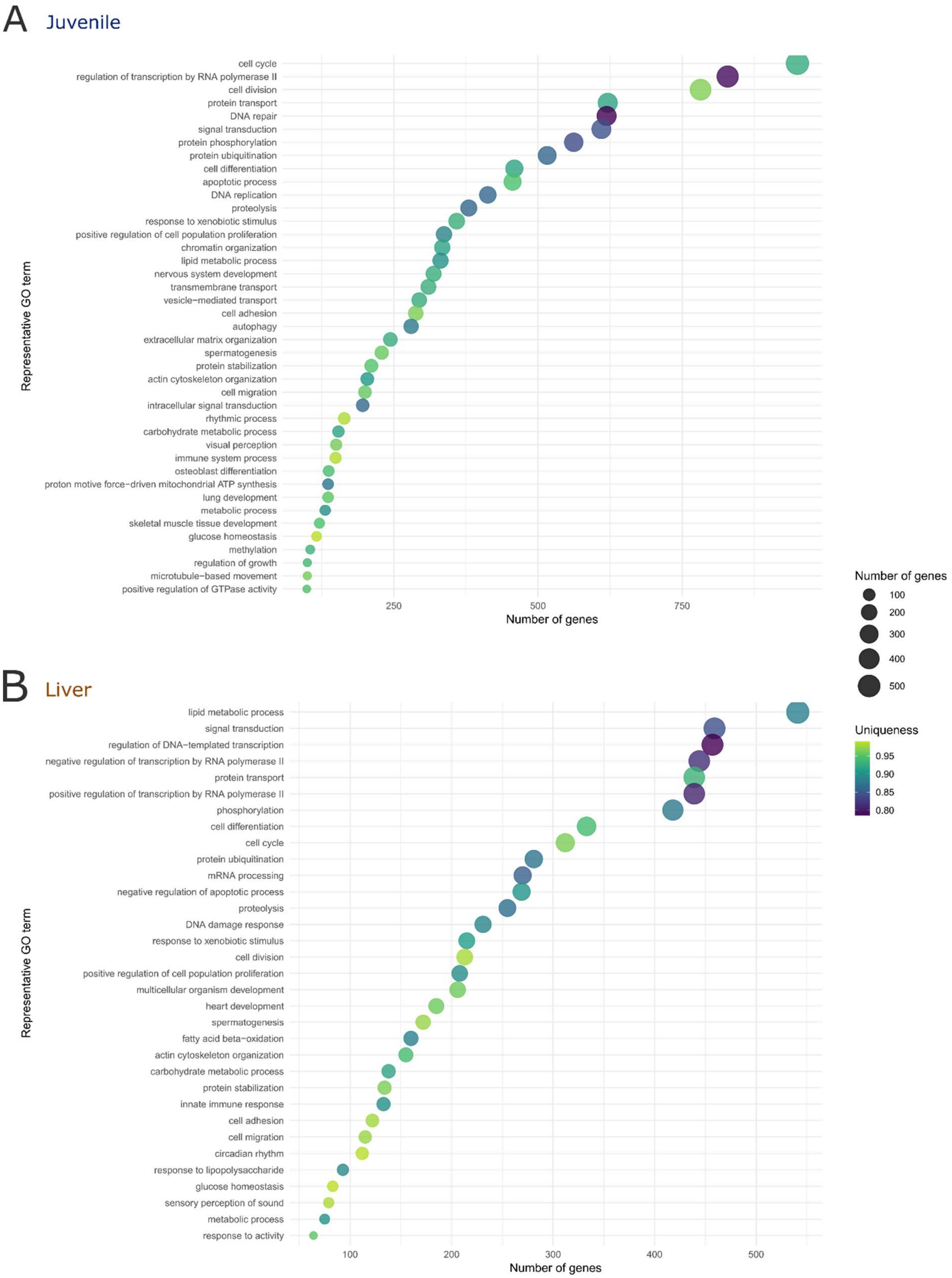


**Extended Data Fig. 3: Most common GO terms across the top 10 percent genes driving the loading of dimension 1 on the Principal Component Analysis (PCAs) of the feeding experiment samples for juveniles (a) and adult liver (b).** The PCAs are presented on Fig. 3. GO terms were trimmed based on semantic similarity at a threshold of 0.7 using GOSemSim and the most unique terms are presented here.

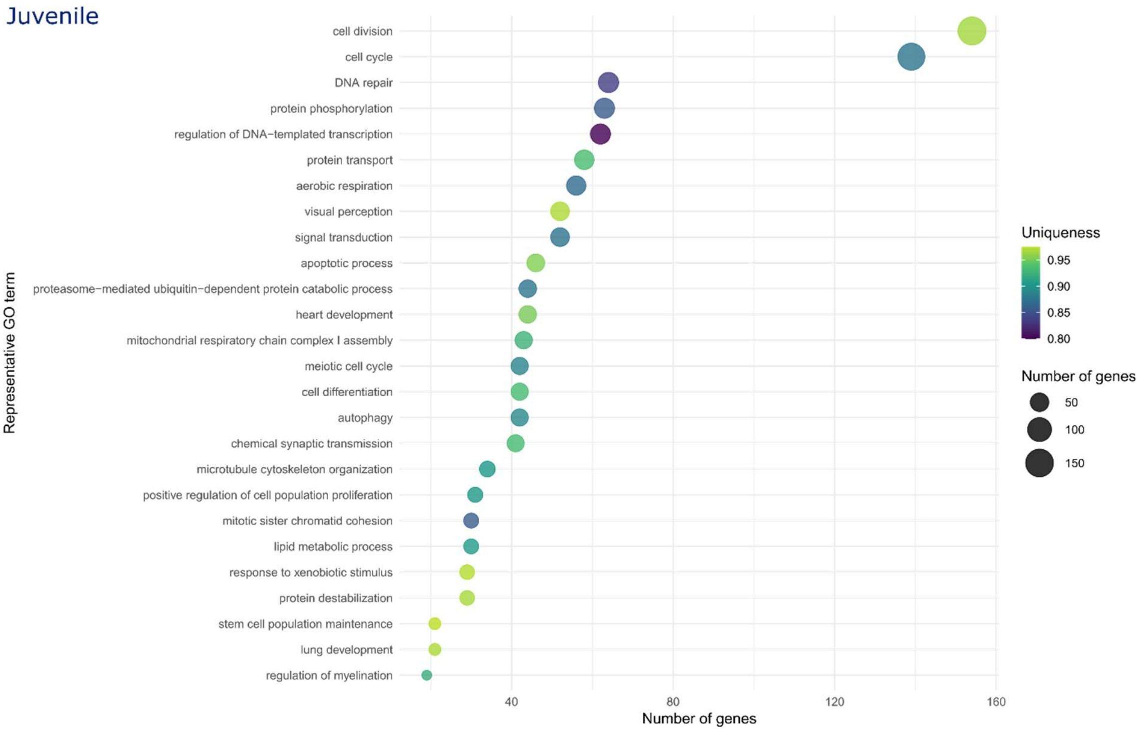


**Extended Data Fig. 4: Most common GO terms across the top 10 percent genes driving the loading of dimension 2 on the Principal Component Analysis of the feeding and field samples (Fig. 3b from the main text) in juveniles.** GO terms were trimmed based on semantic similarity at a threshold of 0.7 using GOSemSim and the most unique terms are presented here.

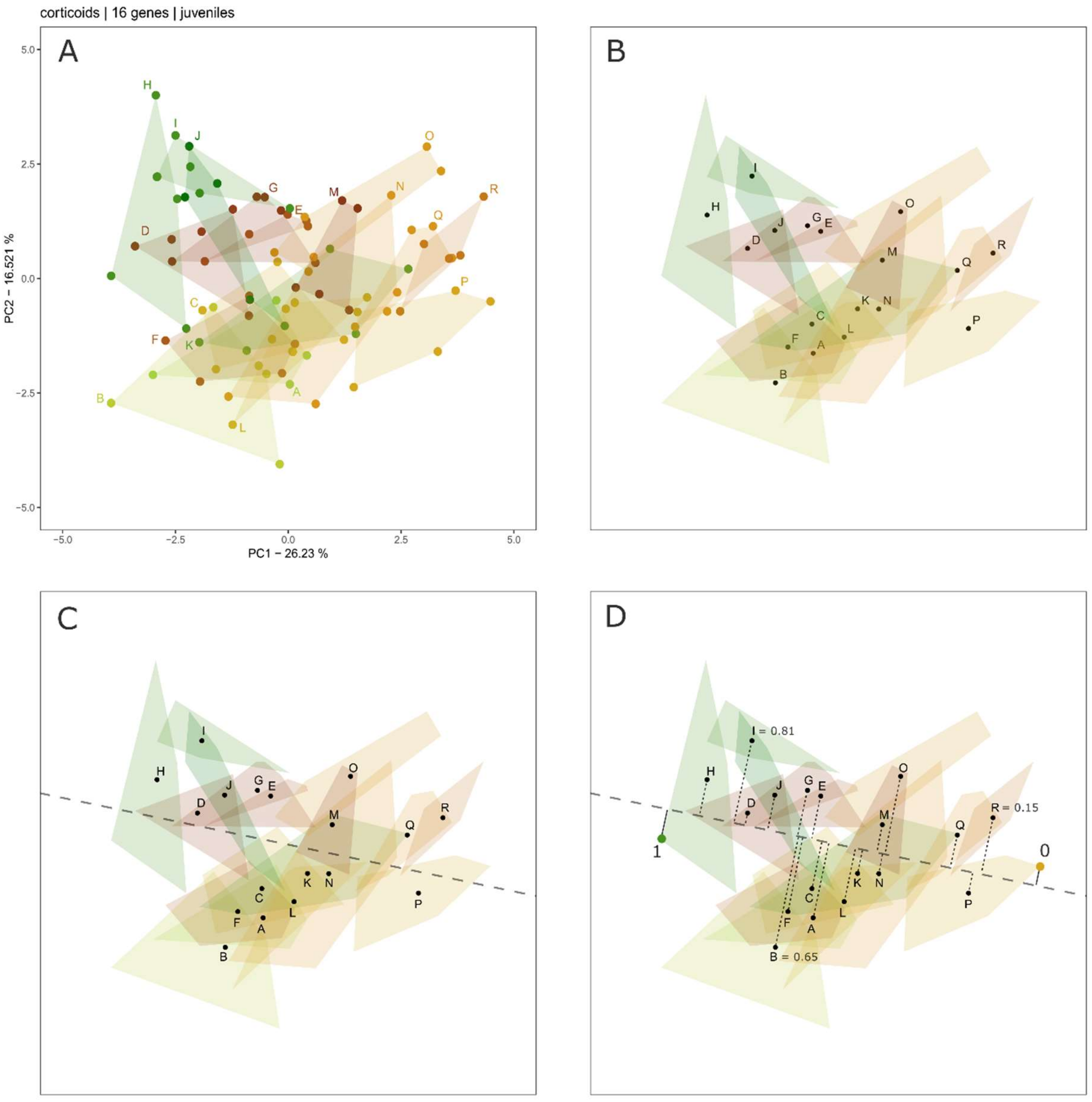


**Extended Data Fig. 5: Sketch of the method to obtain the radar plots presented in Fig. 4.** This is an example where the first two PC dimensions are used. The analysis includes all PC dimensions representing up to 50% of the total variance. **a**, First, the Principal Component Analysis, here based on the normalized expression level of 16 genes linked to corticoids, across 18 sites in Okinawa, with 5 samples per site, is obtained. **b**, The mean PC1/PC2 coordinate of the samples from each of the 18 sites is calculated. **c**, The direction of the main variance in the mean coordinate of the sites is then calculated, here presented as a dashed axis. **d**, The orthogonal projection of all mean site coordinates as well as the projection of each sample onto the main axis of variance is calculated. The two furthermost samples across the main variance axis are set to have values of 0 and 1. Based on this method, the orthogonal projection of all samples and sites was calculated for multiple gene sets. The results are then summarized as a radar plot in the main manuscript.

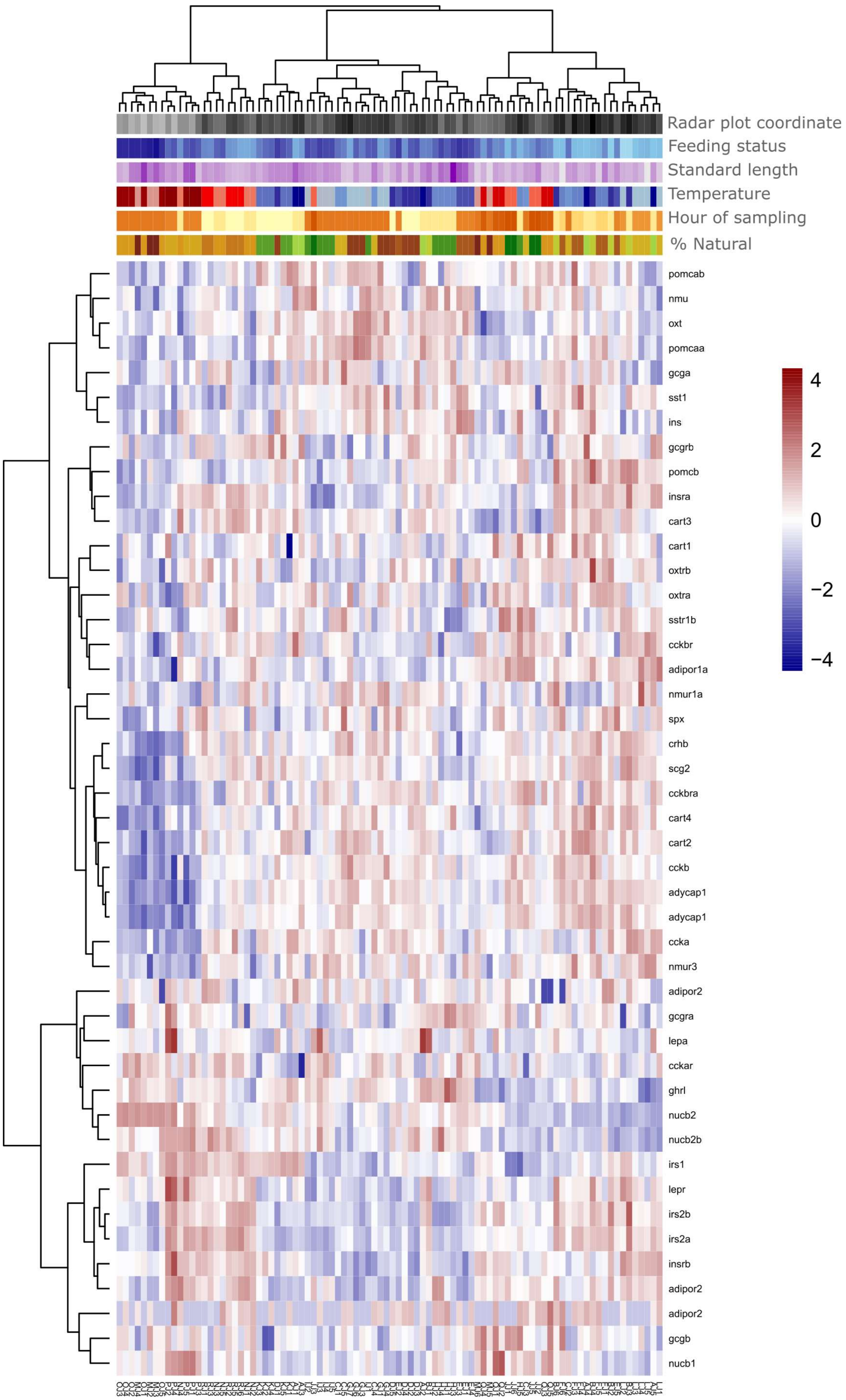
Radar plot coordinate
Feeding status
Standard length
Temperature
Hour of sampling
% Natural
pomcab
nmu
oxt
pomcaa
gcga
sst1
ins
gcgrb
pomcb
insra
cart3
cart1
oxtrb
oxtra
sstr1b
cckbr
adipor1a
nmur1a
spx
crhb
scg2
cckbra
cart4
cart2
cckb
adycap1
adycap1
ccka
nmur3
adipor2
gcgra
lepa
cckar
ghrl
nucb2
nucb2b
irs1
lepr
irs2b
irs2a
insrb
adipor2
adipor2
gcgb
nucb1
4
2
0
−2
−4

**Extended Data Fig. 6: Heatmap of the vsd-normalised expression level of 45 appetite-related genes in juvenile field samples.** Rows are clustered based on correlation using the Ward D2 method with *pheatmap*. Environmental (temperature, hour of sampling) and intrinsic fish variables (feeding status, standard length) are indicated above the heatmap. Increasingly dark colours indicate higher values for the radar plot coordinates, feeding status, standard length, and hour of sampling. Temperature is indicated from a blue (low temperature) to red (high temperature) scale.

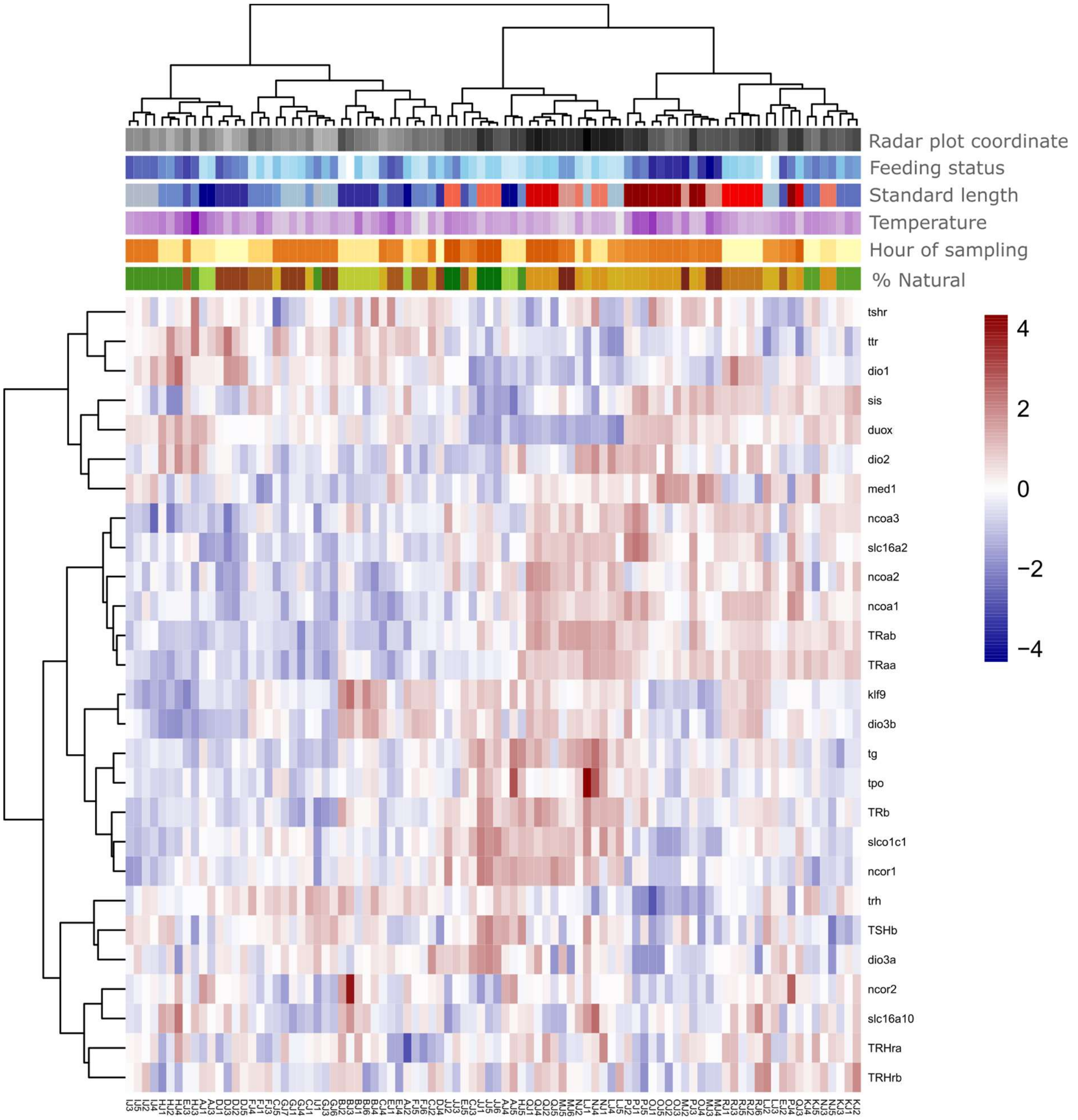


**Extended Data Fig. 7: Heatmap of the vsd-normalised expression level of 27 thyroid-related genes in juvenile field samples.** Rows are clustered based on correlation using the Ward D2 method with *pheatmap*. Environmental (temperature, hour of sampling) and intrinsic fish variables (feeding status, standard length) are indicated above the heatmap. Increasingly dark colours indicate higher values for the radar plot coordinates, feeding status, standard length, and hour of sampling. Temperature is indicated from a blue (low temperature) to red (high temperature) scale.

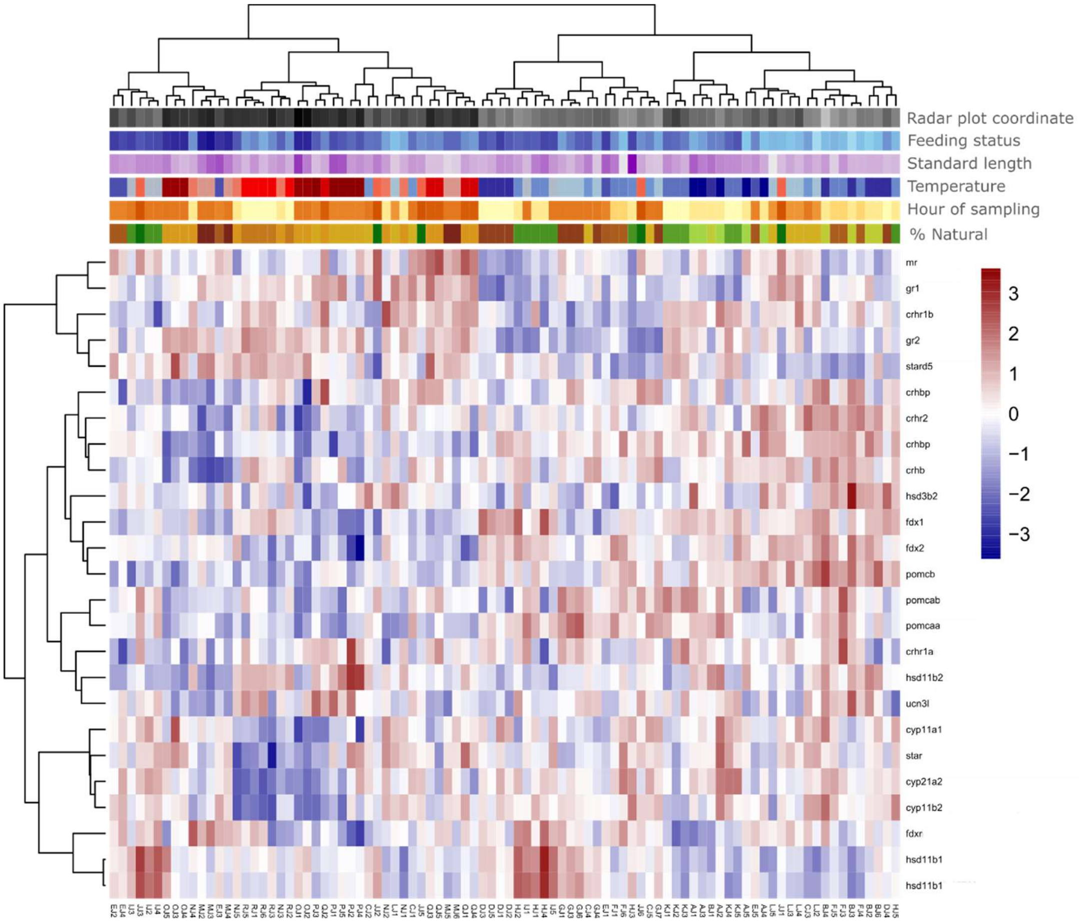


**Extended Data Fig. 8: Heatmap of the vsd-normalised expression level of 25 corticoid-related genes in juvenile field samples**. Rows are clustered based on correlation using the Ward D2 method with *pheatmap*. Environmental (temperature, hour of sampling) and intrinsic fish variables (feeding status, standard length) are indicated above the heatmap. Increasingly dark colours indicate higher values for the radar plot coordinates, feeding status, standard length, and hour of sampling. Temperature is indicated from a blue (low temperature) to red (high temperature) scale.

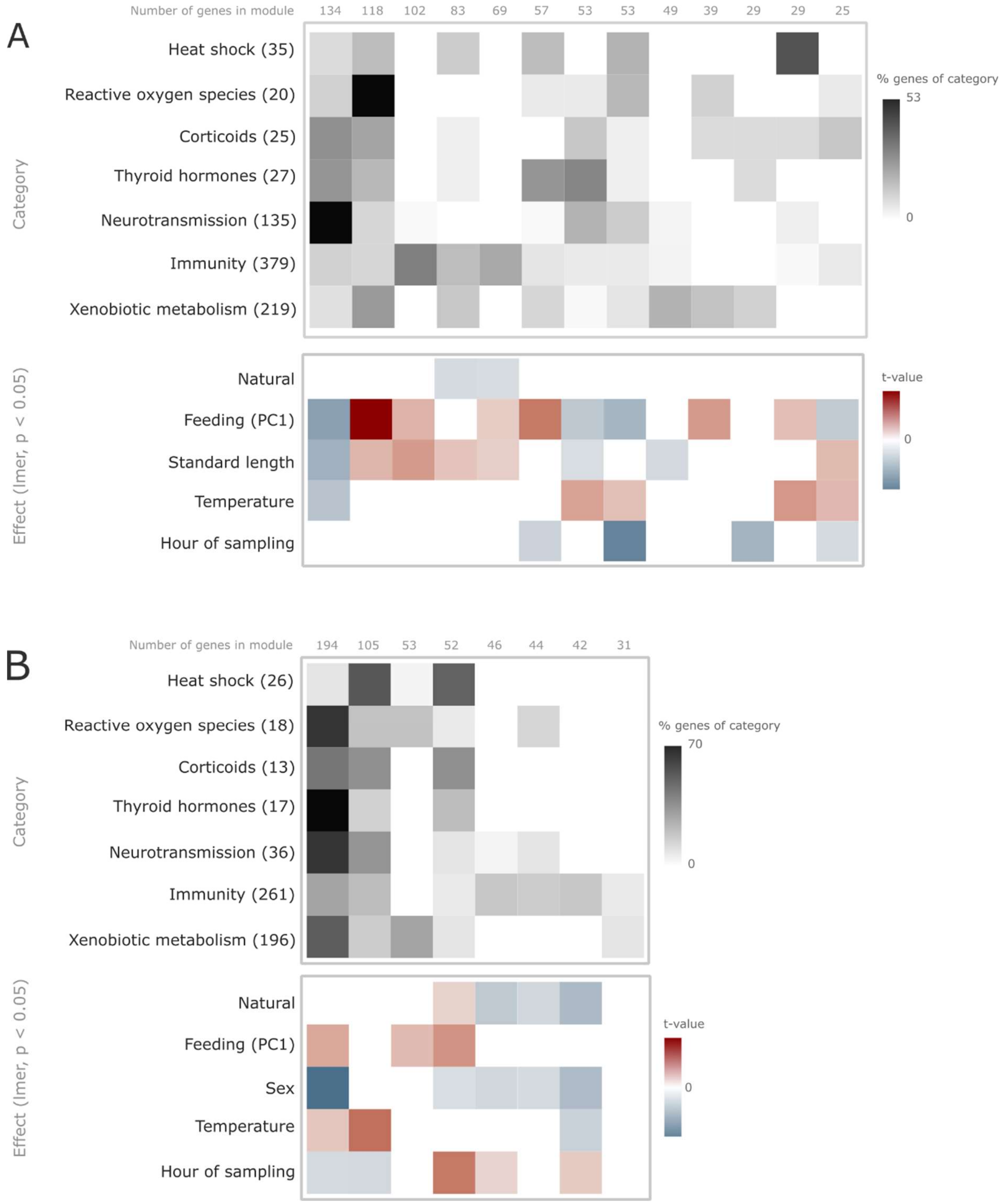


**Extended Data Fig. 9: Results from a module analysis for the juvenile whole-body and adult liver gene expression.** Analysis performed using WGCNA (Weighted Correlation Network Analysis) with a minimum module size of 20 genes for juveniles and 13 genes for adult liver samples (corresponding to the lowest number of genes within a category) and a soft-thresholding power of 9 for juveniles and 6 for livers. 13 modules of genes with similar expression patterns across the samples were retrieved for the juveniles and 8 for the livers. The following categories of genes were used, with 974 genes retrieved in total: heat shock proteins, reactive oxygen species, corticoids, thyroid hormones, neurotransmission, immunity, and xenobiotic metabolism. Environmental and intrinsic fish variables (percentage of natural zones within a kilometre of the site, feeding status as derived from Fig. 3 of the main text, standard length for juveniles, sex for livers, temperature, and hour of sampling) were used in a linear-mixed effect model (with site as a random effect and a selection of fixed effects based on model dredging) to determine the relationship between the gene expression patterns in each module and environmental or intrinsic fish drivers. The resulting t-values are presented. Higher t-values (in red) indicate positive relationships between the gene expression level within the module and the given variable, and inversely (in blue). Top: Percentage of genes of each category retrieved within a given module. Bottom: t-value of a linear mixed-effect model retrieving significant relationships between environmental or intrinsic fish characteristics and gene expression patterns within the modules.

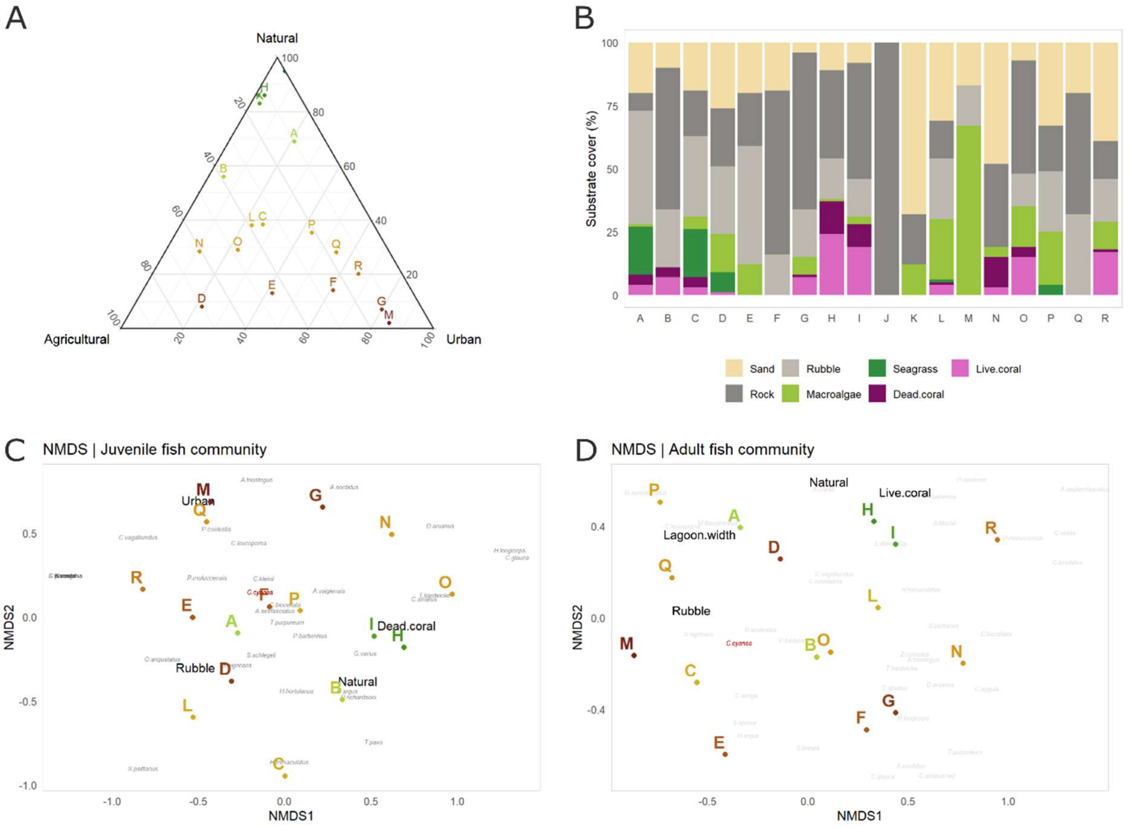


**Extended Data Fig. 10: Environmental characterisation of the sites. a,** Ternary plot presenting the percentage of natural, agricultural, and urban zones within 1km of the sites (excluding water). **b,** Substrate cover on three 25m transects performed parallel to shore near the sampling (maximum depth: 2m). **c,** NMDS analysis of the juvenile fish community structure and contributing environmental variables based on *vegan* function envfit ($p < 0.1$). **d,** NMDS analysis of the adult fish community structure and contributing environmental variables based on *vegan* function envfit ($p < 0.1$). No fish community structure was recorded on sites J and K due to logistical constraints.

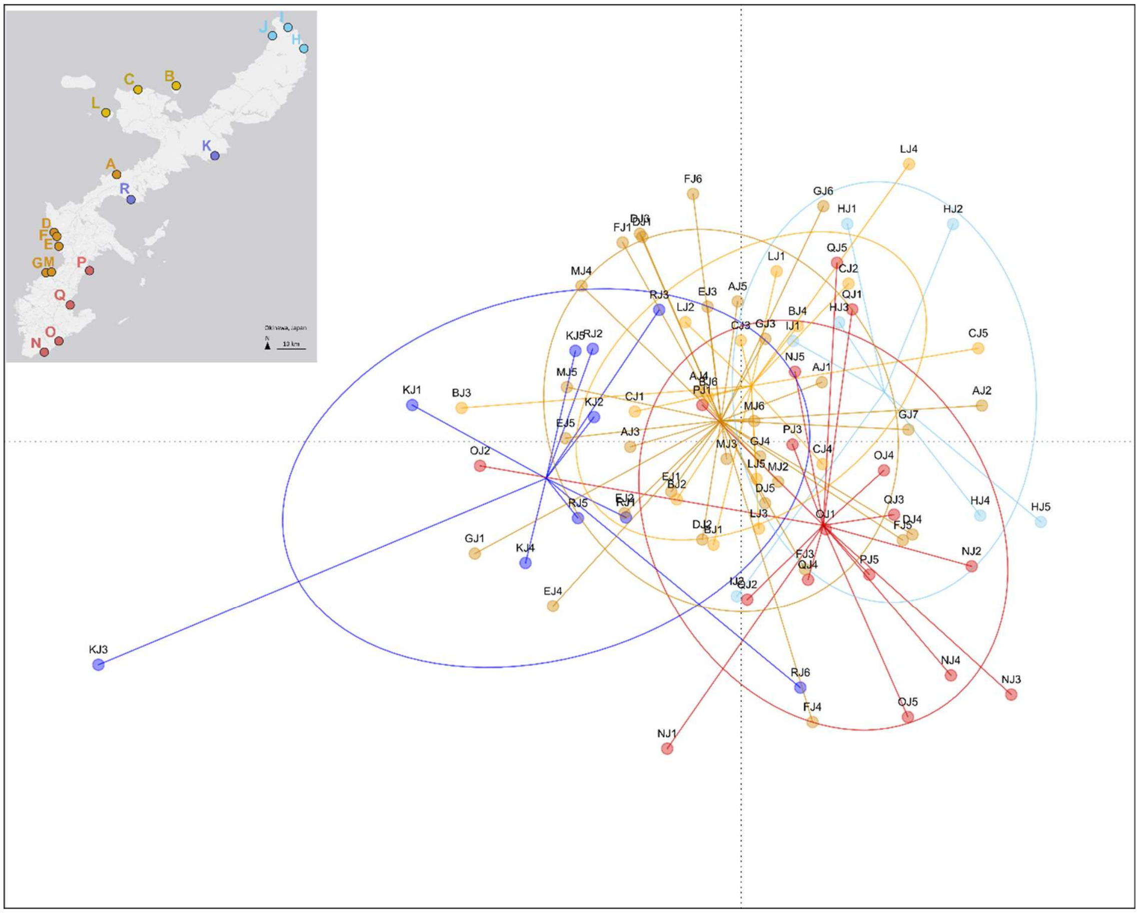


**Extended Data Fig. 11: First two discriminant functions of a Discriminant Analysis of Principal Components (DAPC) of *C. cyanea* juveniles.** Colours indicate 5 zones: northwest, north, east, south, west. The analysis is based on filtered SNPs (mind = 0.25, geno = 0.25) following variant calling using the Genome Analysis Toolkit (GATK) pipeline.
